# Spatial analysis of IPMNs defines a paradoxical KRT17-positive, low-grade epithelial population harboring malignant features

**DOI:** 10.1101/2025.03.18.643943

**Authors:** Jay Li, Georgina Branch, Justin Macchia, Ahmed M. Elhossiny, Nandini Arya, Julia Liang, Padma Kadiyala, Nicole Peterson, Richard Kwon, Jorge D. Machicado, Erik-Jan Wamsteker, Allison Schulman, George Philips, Stacy Menees, Jonathan Xia, Aatur D. Singhi, Vaibhav Sahai, Jiayun M. Fang, Timothy L. Frankel, Filip Bednar, Marina Pasca di Magliano, Jiaqi Shi, Eileen S. Carpenter

## Abstract

**Background & Aims:** Intraductal papillary mucinous neoplasms (IPMNs) are pancreatic cysts that represent one of the few radiologically identifiable precursors to pancreatic ductal adenocarcinoma (PDAC).

Though the IPMN-bearing patient population represents a unique opportunity for early detection and interception, current guidelines provide insufficient accuracy in determining which patients should undergo resection versus surveillance, resulting in a sizable fraction of resected IPMNs only harboring low-grade dysplasia, suggesting that there may be overtreatment of this clinical entity.

**Methods:** To investigate the transcriptional changes that occur during IPMN progression, we performed spatial transcriptomics using the Nanostring GeoMx on patient samples containing the entire spectrum of IPMN disease including low-grade dysplasia, high-grade dysplasia, and IPMN-derived carcinoma. Single cell RNA sequencing was performed on side branch and main duct IPMN biospecimens.

**Results:** We identified a subpopulation of histologically low-grade IPMN epithelial cells that express malignant transcriptional features including *KRT17*, *S100A10* and *CEACAM5*, markers that are enriched in PDAC. We validated this high-risk gene signature in both single-cell RNA sequenced samples and an external ST dataset containing a larger number of IPMN samples including non-tumor bearing IPMN (i.e. low-grade IPMN in isolation). Immunofluorescence staining of a large cohort of patient tissues confirmed the presence of KRT17-positive cells, which were found to comprise a small subset of epithelial cells within histologically low-grade IPMN in a patchy distribution.

**Conclusions:** Our study demonstrates that KRT17 marks a distinct transcriptional signature in a subpopulation of epithelial cells within histologically low-grade IPMN. This population of cells likely represents a transitional state of histologically low-grade epithelial cells undergoing progression to a higher grade of dysplasia and thus may represent a higher risk of progression to carcinoma.

Graphical Abstract

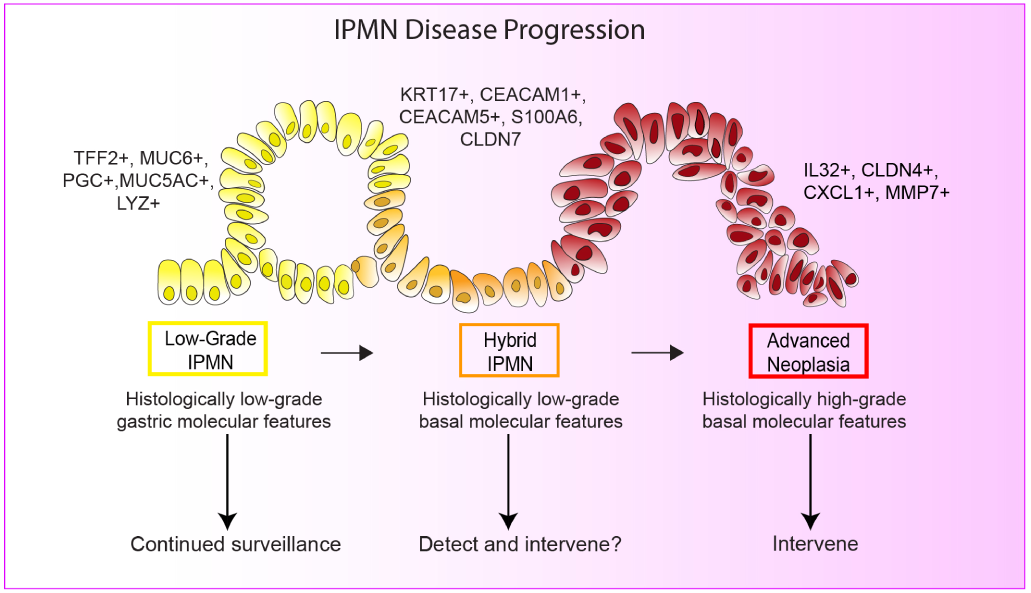

## Introduction

Pancreatic ductal adenocarcinoma (PDAC) has a very poor prognosis with a 5-year survival of 13% [1]. It is projected to become the 2^nd^ leading cause of cancer death in the United States by 2040[2]. A significant contributor to poor survival is the observation that the vast majority of PDAC patients are diagnosed with advanced, unresectable disease.

While PDAC is one of the leading causes of cancer death in the US, the incidence of PDAC is quite low, which complicates the development of effective screening tests and early detection methods. While most PDAC cases arise from pancreatic intraepithelial neoplasms (PanIN), up to 15% of PDAC cases arise from mucinous pancreatic cysts, most of which are intraductal papillary mucinous neoplasms (IPMNs) [3]. Mucinous pancreatic cysts are the only radiologically identifiable precursor to PDAC. The diagnosis, often incidental, of these lesions is rising with the growing use of abdominal cross-sectional imaging such as computed tomography (CT) and magnetic resonance imaging (MRI)[4]. Thus, the patient population with IPMN is a relatively identifiable subset of individuals and represent a unique opportunity for early detection of PDAC.

IPMNs are classified based on degree of dysplasia and histologic subtype (gastric, intestinal, pancreaticobiliary) [5, 6]. These cystic lesions may undergo traditional histologic progression from low-grade to high-grade dysplasia to invasive carcinoma. KRAS and GNAS mutations are common across histologic subtypes with additional mutations in RNF43 frequent in the intestinal subtype and TP53 and SMAD4 mutations in the pancreaticobiliary subtype [7–10]. While IPMNs are quite prevalent, being detected in up to 10% of the population above age 50, only a small proportion of IPMNs progress to PDAC [11]. The two histological entities that should prompt surgical resection are the presence of high-grade dysplasia or carcinoma, both of which are categorized as advanced neoplasia in IPMN. Unfortunately, the only method to accurately determine the presence of advanced neoplasia is histopathological analysis of the entire cyst, which can only be achieved in the setting of an invasive surgical resection. There have been several clinical guidelines that attempt to risk-stratify patients based on a combination of non-invasive methods, including imaging findings, cyst fluid markers and cytology, and clinical symptoms [12, 13]. However, current methodologies have insufficient specificity in identifying advanced neoplasia[14]. As a result, prior studies have identified a sizable fraction of resected IPMNs only harboring low-grade dysplasia, suggesting that there may be overtreatment of this clinical entity for which the recommended treatment is surgery with high potential for morbidity and mortality [15, 16]. A major gap in the field is the lack of effective strategies to accurately risk-stratify pancreatic cysts and determine which IPMNs are at higher risk of progression.

To define the transcriptional changes underlying IPMN progression, we utilized a spatial transcriptomic (ST) profiling approach using the NanoString GeoMx Digital Spatial Profiler platform on archival patient tissue containing the entire spectrum of IPMN disease including low-grade IPMN (LG-IPMN), high-grade IPMN (HG-IPMN), and IPMN-derived PDAC from surgically resected IPMN samples. This approach provides advantages over bulk or single-cell RNA sequencing including preservation of spatial relationships and correlation of extracted RNA from each selected region of interest (ROI) with gold standard pathologic diagnosis.

Here, we identify a subset of LG-IPMN that, despite being histologically low-grade, exhibits transcriptomic similarities to HG-IPMN and PDAC. The presence of this paradoxical “hybrid” low-grade population suggests that malignant transcriptional alterations are active prior to histologic progression and may be leveraged to identify patients with higher risk disease. We next extracted the transcriptomic signatures of this higher risk LG-IPMN subgroup to query in both our single-cell RNA sequencing dataset of IPMN samples as well as a larger published ST dataset including non-tumor bearing LG-IPMN. Finally, we validated the presence of these hybrid cells utilizing patient pancreatic cyst tissue microarrays, demonstrating that these cells, while not pervasive throughout the cyst epithelium, are present in LG-IPMN in a small proportion. Altogether, these analyses shed light on the changes that may occur as benign pancreatic cysts progress, and provide potential targets for development of early detection tools and predictive biomarkers to improve risk-stratification for pancreatic cysts.

## Results

### LG-IPMNs display transcriptional heterogeneity

To investigate the transcriptomic changes underlying histologic progression of IPMNs, we utilized the NanoString GeoMx Digital Spatial Profiler platform to perform whole transcriptome profiling on formalin-fixed paraffin-embedded tissue sections from surgical resections of 3 patients with treatment-naïve IPMN with associated invasive carcinoma **(Supplemental Figure 1A)**. Utilizing expert clinical pathologist annotation, ROIs corresponding to acini, normal duct, LG-IPMN, HG-IPMN, and tumor were selected (**Figure 1A**). User-selected epithelial regions-of-interest (ROIs) were segmented by PanCK expression to enrich for transcripts from epithelial populations, resulting in 12 acinar ROIs, 13 normal duct ROIs, 83 LG-IPMN ROIs, 54 HG-IPMN ROIs, and 16 tumor ROIs after pre-processing and filtration (**Figure 1B**). To determine how the different histologically defined cell types related based on transcriptional features, we calculated the top variable genes (with a coefficient of variation in the top 20%) for each ROI and performed hierarchical clustering of the epithelial ROIs based on these genes. Histologically benign epithelial ROIs such as acinar, normal duct, and the majority of LG-IPMN clustered together, while HG-IPMN and tumor clustered together, as expected (**Figure 1C**). Surprisingly, we observed that a small subset of LG-IPMN ROIs clustered with HG-IPMN and tumor (**Figure 1C, blue clade and box)** This was further visualized on principal component analysis (PCA) plot, which showed a heterogeneous spectrum of LG-IPMN ranging from ROIs overlapping primarily with normal ducts to those overlapping with HG-IPMN and tumor (**Figure 1D, Supplemental Figure 1B**). We wished to further study these regions that had divergent histologic and transcriptomic features and thus reclassified this subset of LG-IPMN ROIs to henceforth refer to them as “hybrid IPMN” (Hyb-IPMN) (**Figure 1D**). On careful review of histology, no apparent morphological features could readily distinguish between the Hyb-IPMN ROIs and LG-IPMN ROIs (**Supplemental Figure 2).** These findings highlight the transcriptional heterogeneity within LG-IPMNs and suggest the existence of a distinct subset that shares molecular features with more advanced disease stages.

**Figure 1:**
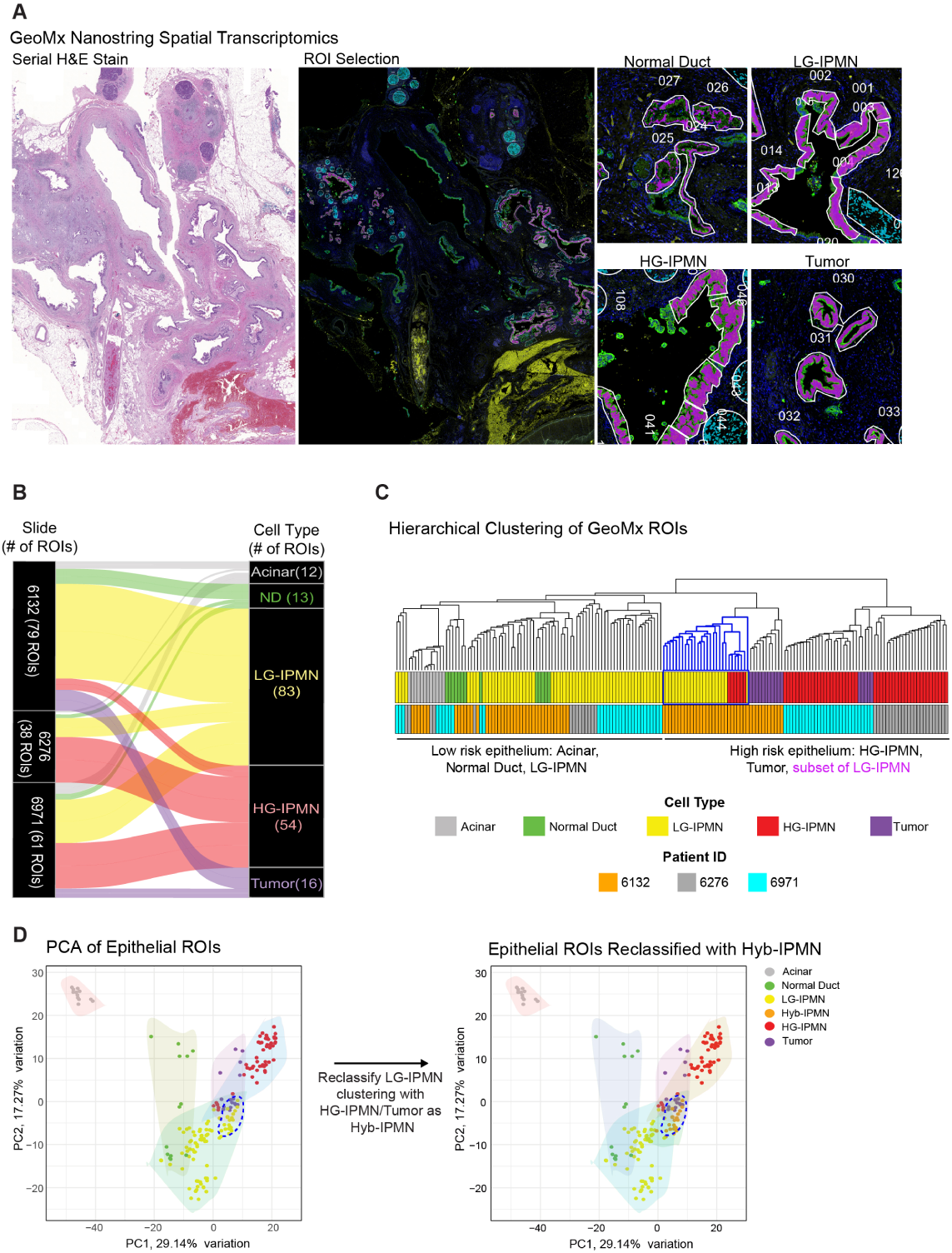
Spatial transcriptomics reveals transcriptional heterogeneity in low-grade IPMN. **A)** GeoMX tissue section with corresponding serially sectioned H&E stain from a surgically resected IPMN with associated carcinoma, panCK and CD45. panCK^+^ segments are pseudocolored in green, whereas panCK^−^CD45^−^ segments are pseudocolored in yellow. Purple pseudocolor represents the PanCK segmented cells selected in the user ROIs. **B)** Sankey plot showing the distribution of ROIs in IPMN samples. **C)** Hierarchical clustering of ROIs based on variably expressed genes. Blue clade and box marks ROIs that clustered together despite ranging from LG-IPMN to HG-IPMN histologically. **D)** UMAP overlay of disease states on extracted epithelial cells from the single-cell dataset of healthy, adjacent normal, and tumor samples. **E)** PCA plot of epithelial segmented ROIs (each dot represents one ROI) before and after reclassification of LG-IPMN ROIs that clustered with HG-IPMN/Tumor ROIs as Hyb-IPMN.

### A subpopulation of LG-IPMN epithelium shares high-risk transcriptomic features with advanced neoplasia

While direct transcriptomic comparisons between LG-IPMN, HG-IPMN, and IPMN-derived tumor have been performed at the bulk, single-cell, and spatial level [17–20], the heterogeneity of gene expression within histologically low-grade appearing epithelium has been less explored. Thus, we focused on the features distinguishing Hyb-IPMN from LG-IPMN and performed differential expression analysis to identify differentially expressed genes (DEGs) in LG-IPMN versus Hyb-IPMN, thus identifying 43 DEGs enriched in LG-IPMN and 73 DEGs enriched in Hyb-IPMN (**Figure 2A, Supplementary Tables 1** **and 2**).

**Figure 2:**
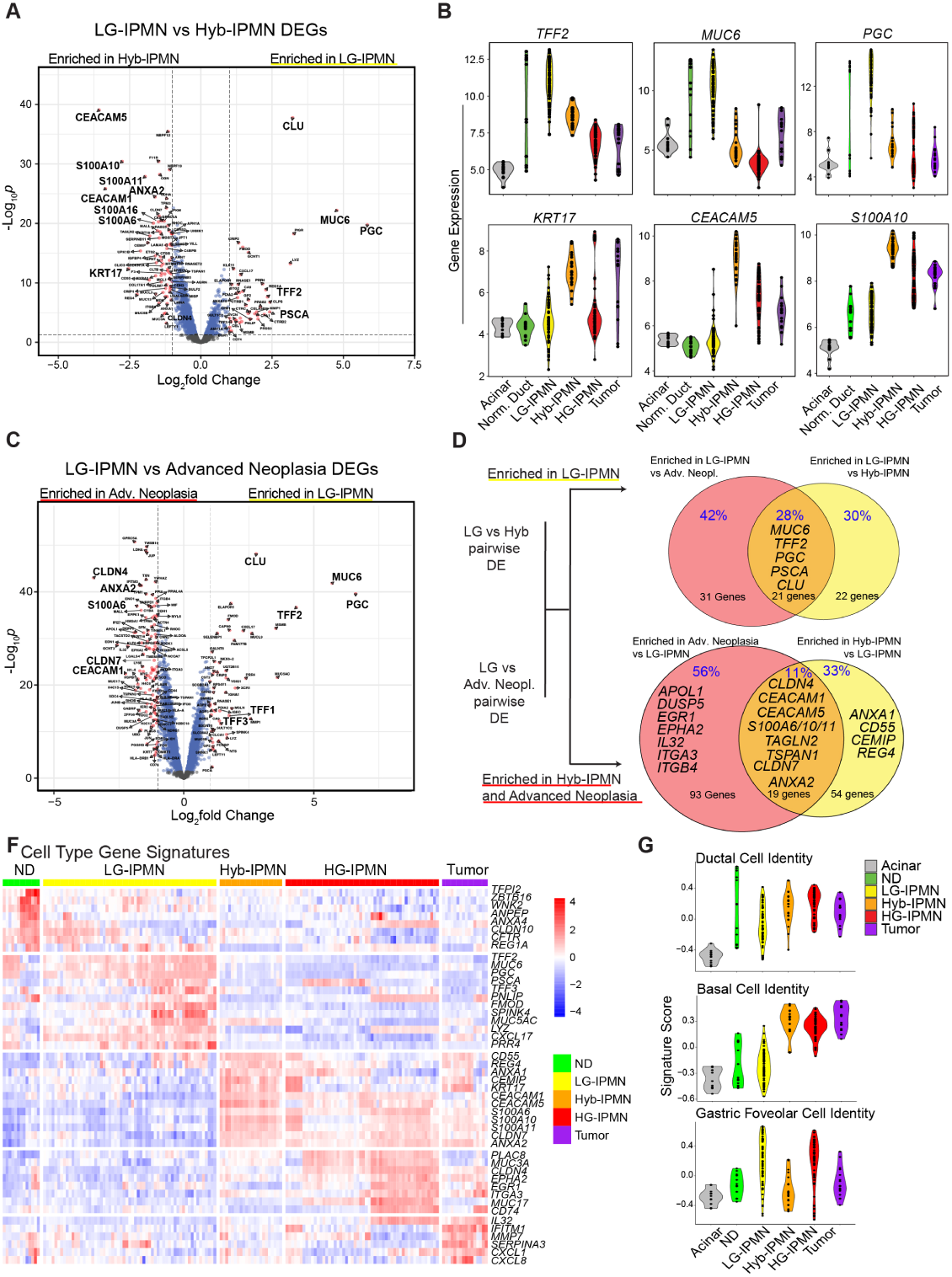
Hyb-IPMN shares high-risk transcriptomic features with advanced neoplasia. **A)** Volcano plot of all DEG (Differentially Expressed Genes) comparing Hyb-IPMN to LG-IPMN. Genes that are significantly differentially expressed, as determined by adjusted P value of <0.05, are colored in red. **B)** Violin plots of gene expression in each ROI, grouped by cell type. **C)** Volcano plot of all DEG (Differentially Expressed Genes) comparing Advanced Neoplasia (consisting of HG-IPMN and carcinoma) to LG-IPMN. Genes that are significantly differentially expressed, as determined by adjusted P value of <0.05, are colored in red. **D)** Venn Diagrams showing shared genes that are enriched in advanced neoplasia and Hyb-IPMN compared to LG-IPMN (top) and enriched in LG-IPMN compared to advanced neoplasia and Hyb-IPMN (bottom). **F)** Heatmap of cell type–specific markers derived from linear mixed model differential gene expression on ROIs. **G)** Violin plots of the GSVA scores of Panglao cell identity signatures for ductal cell, basal cell, and gastric foveolar cell identity on ROIs.

The genes most enriched in LG-IPMN compared to Hyb-IPMN included primarily gastric genes such as PGC, MUC6, and TFF2, an expected finding as they have been previously reported markers of LG-IPMN (**Figure 2B**) [21–24]. Genes enriched in Hyb-IPMN included known PDAC-associated genes such as *CEACAM1, CEACAM5, KRT17, CLDN4, S100A6, S100A10,* and *S100A11* [25–31]. Among these, the top 4 enriched transcripts in Hyb-IPMN were *CEACAM5, CEACAM1, S100A10,* and *KRT17* (**Figure 2B**), with the latter a marker of the more aggressive basal subtype of PDAC that is associated with poor prognosis [26, 32, 33]. Given the enrichment of PDAC-associated transcripts in Hyb-IPMN, we hypothesized that these ROIs represent a distinct group of epithelial cells with shared transcriptional features of both low-grade dysplasia and carcinoma, and thus potentially are at higher risk of progression to invasive disease.

To further investigate the notion that Hyb-IPMN represents a transitional state between LG-IPMN and advanced neoplasia (representing the combination of HG-IPMN and tumor), we performed differential expression analysis between LG-IPMN and advanced neoplasia ROIs, (**Figure 2C)**, then queried genes that are similarly enriched in both Hyb-IPMN and advanced neoplasia compared to LG-IPMN (**Figure 2D**). These overlapping DEGs represent expression changes that precede histologic progression and are retained in advanced neoplasia, suggesting a potential role in early IPMN pathogenesis. LG-IPMN overlapping genes included the previously identified gastric genes *PGC, MUC6*, and *TFF2* (**Figure 2D**). Thus, the downregulation of these gastric genes may represent an early event in the progression of LG-IPMN that occurs prior to histologic progression, as LG-IPMN and Hyb-IPMN are histologically indistinguishable. We next queried DEGs that were upregulated in both Hyb-IPMN and advanced neoplasia compared to LG-IPMN (**Figure 2D**). In the list of 19 overlapping DEGs, we noted several shared enriched genes upregulated in and associated with poor prognosis in PDAC, including *CEACAM1, CEACAM5, CLDN4, S100A6, S100A10,* and *S100A11* [27–31].

Plotting gene expression of top-expressing low-grade and Hyb-IPMN markers revealed nearly mutually exclusive expression between LG-IPMN and advanced neoplasia (HG-IPMN and tumor), while Hyb-IPMN shared several features with advanced neoplasia (**Figure 2F, Supplementary Table 3)**. Consistent with the hierarchical clustering and PCA (**Figure 1C-D**), the Hyb-IPMN transcriptome harbored many features which overlapped with LG-IPMN or tumor ROI transcriptomes. Gene set variation analysis revealed that while upregulation of apical junction genes was shared in Hyb- and HG-IPMN compared to LG-IPMN and largely driven by CLDN4/7, upregulation of inflammatory pathways in TNFα signaling and IFNγ response was limited to HG-IPMN (**Supplemental Figure 3A-C)**. As recent studies have demonstrated that *NKX6-2*, a gastric transcription factor, drives gastric transcriptional programs in LG-IPMN and is linked to a more indolent disease course [19, 34], we performed gene signature scoring on the ROIs for cell identity utilizing the Panglao database, which contains curated gene signatures for various cell types based on single-cell RNA sequencing data[35]. We found that while normal ducts, LG-, Hyb-, and HG-IPMNs, and tumors have similar scores in ductal cell identity, the basal cell identity was elevated in Hyb-IPMN, HG-IPMN, and tumor ROIs, consistent with the acquisition of a basal gene program promoting progression to malignancy (**Figure 2G)**.

Interestingly, there was heterogeneous enrichment of the gastric foveolar signature in both LG-IPMN and HG-IPMN, driven by genes including *PGC, TFF2 and GKN2*. However, the gastric foveolar signature was uniformly reduced in normal ducts, Hyb-IPMN, and tumor ROIs, suggesting the notion that Hyb-IPMN may represent a “branch-point” in the trajectory of malignant progression of IPMNs.

### Hyb-IPMN shares features with both PanIN and PanIN-derived carcinoma

We have previously found that low-grade PanIN is highly prevalent in the general population; to determine whether our Hyb-IPMN signature could give us insights on the evolution of PanIN to carcinoma, we queried our published dataset, which included of PanIN from healthy donor pancreata and PanIN-derived carcinoma from patient PDAC tissues[36]. We found that LG-IPMN were transcriptomically most similar to PanINs from healthy donors who had no underlying pancreatic disease **(Figure 3A)**. HG-IPMN were most correlated with glandular (well-differentiated) PDAC, while Hyb-IPMN interestingly were also most correlated with Glandular PDAC, although to a lesser degree **(Figure 3A)**. In comparing specific upregulated genes between cell types, we found that *MUC5AC, S100A6/A10* and *TFF1/2* were common in Hyb-IPMN and both PanIN from healthy and tumor-bearing pancreas, however, *KRT17* upregulation in pre-cancerous lesions was exclusive to Hyb-IPMN **(Figure 3B-C, Supplementary Table 4)**. Additionally, PanIN-driven tumor progression was characterized by a sustained gastric differentiation signature, as evidenced by persistent expression of *PGC, TFF2,* and *MUC6,* which remained detectable even at the stage of glandular PDAC **(Figure3D)**. IPMN-driven progression, in contrast, showed a distinct decrease in these markers in HG-IPMN and Hyb-IPMN, which precede carcinoma **(Figure2B).** These data suggest key differences in the evolution of IPMN-derived carcinoma compared to PanIN-derived carcinoma. First, KRT17 upregulation in IPMNs appears earlier, specifically at the histologic LG-IPMN stage, albeit in only a subset of cells, while in PanIN-derived carcinoma, KRT17 upregulation is only seen in carcinoma. Second, the marked loss of gastric differentiation features during IPMN progression is much less notable in PanIN progression.

**Figure 3:**
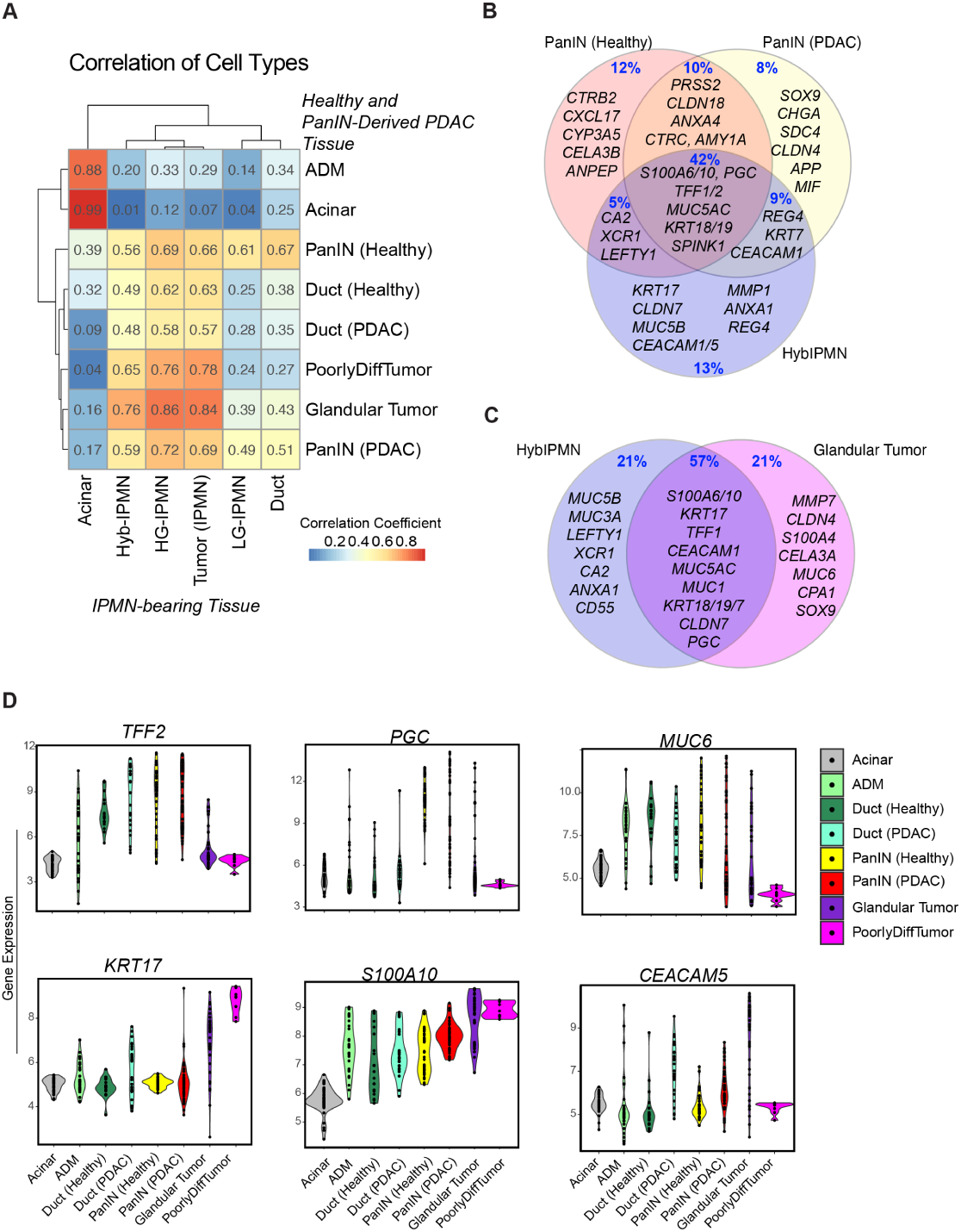
Hyb-IPMN shares features with both PanIN and PanIN-derived carcinoma. A) Correlation plot comparing transcriptional signatures from IPMN and IPMN-derived carcinoma (x-axis) to PanIN and PanIN-derived carcinoma (y-axis). B) Venn Diagram showing shared and exclusive top-expressing genes in PanIN from healthy donors, PanIN from PDAC patients, and Hyb-IPMN. C) Venn Diagram comparing top-expressing genes in Hyb-IPMN and PanIN-derived glandular tumor. D) Violin plots of gene expression in each ROI from PanIN and PanIN-derived carcinoma, grouped by cell type.

### The Hyb-IPMN signature is present at the single cell level and enriched in high-risk IPMN

To assess the presence of our spatial-derived signatures at single cell resolution, we performed single cell sequencing on two freshly acquired IPMN samples. One sample was a side branch IPMN, originating from a branch duct connected to the main pancreatic duct. This sample was obtained via core needle biopsy of a mural nodule within the cyst, performed during endoscopic ultrasound evaluation. Clinical pathological examination of the specimen showed gastric low-grade dysplasia without evidence of advanced neoplasia. The second IPMN specimen was acquired from a total pancreatectomy of a patient with main duct IPMN, where the cyst arises from the main pancreatic duct itself. Given that over half of main duct IPMN progress to malignancy[37], surgical resection is indicated for these cysts. A biospecimen for sequencing was acquired by longitudinally bisecting and shaving off a portion of the dilated main duct.

Clinical pathological examination of the specimen showed multifocal areas of invasive PDAC arising from a background of intestinal IPMN with extensive high-grade dysplasia. For reference comparisons, we integrated these data with our previously published single-cell dataset of PDAC tumors and healthy donor pancreata [36, 38]. As expected, given the sampling strategy above, we captured mostly ductal cells in the IPMN samples **(Figure 4A-B).** Compared to the sidebranch IPMN sample, the main duct IPMN sample was enriched in macrophages to a similar proportion in our PDAC samples, which was not unexpected given that macrophage infiltration is associated with the development of malignancy[39]. IPMN samples had relatively sparse inferred copy number variations, as expected[40], with amplification in chromosome 1 in the sidebranch IPMN sample and amplification in chromosome 7 in the main branch IPMN sample (Supplementary Figure A-B), both of which have been reported previously in in genomic characterization studies of IPMN [41, 42].

**Figure 4:**
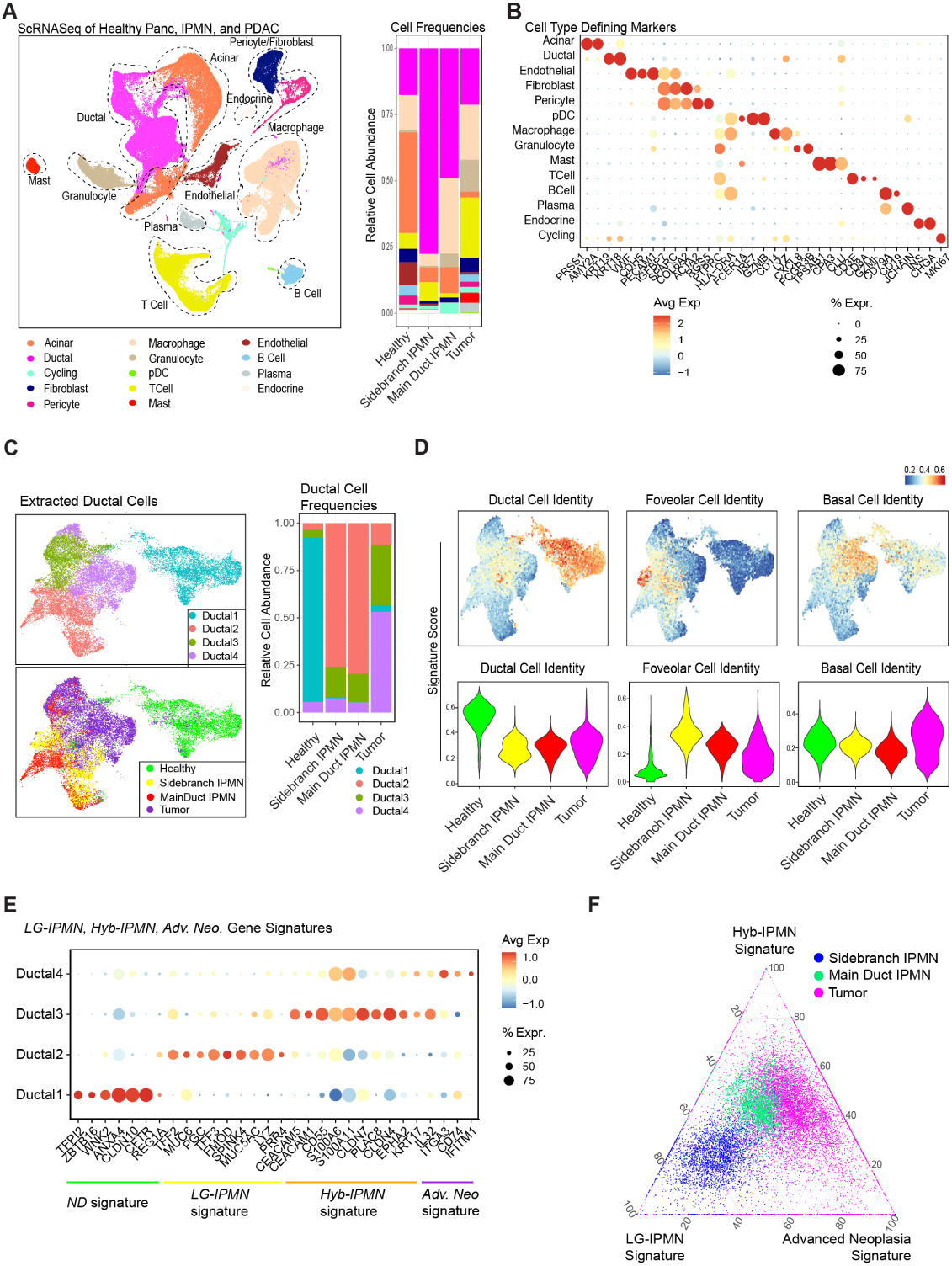
The Hyb-IPMN signature is present at the single cell level and enriched in high-risk IPMN. **A)** Left) UMAP of all cells captured from single-cell RNA sequencing of 2 IPMN, 15 PDAC, and 6 healthy donor pancreas samples. Populations are identified by color. Right) Histogram of cell-type abundance of all cell populations by disease state [healthy, sidebranch IPMN, main duct IPMN, and tumor]. **B)** DotPlot showing markers defining cell populations. **C)** Left) UMAP of epithelial cells from the single-cell dataset of healthy, IPMN, and tumor samples. Populations are identified by color. Right) Histogram of cell-type abundance of all cell populations by disease state [healthy, sidebranch IPMN, main duct IPMN, and tumor].

Extracting the ductal epithelial cells, we obtained 4 specific transcriptionally-defined clusters (**Figure 4C).** The Ductal 1 cluster consisted of normal duct cells, and was enriched in healthy samples specifically, while the Ductal 2 cluster largely consisted of the ductal cells from the side branch and main duct IPMN samples. Ductal 3 and Ductal 4 mostly consisted of malignant epithelial cells from our tumor samples, however there was some overlap with a portion of cells from the main duct IPMN sample, and to a lesser degree the side branch IPMN sample. Scoring each cell for ductal cell, gastric foveolar cell, and basal cell identity, we found that the gastric foveolar identity was largely enriched in Ductal 2 while the basal identity is enriched in Ductal 3/4, in agreement with our spatial data and correlating to the origin tissue type (**Figure 4D**).

Mapping the GeoMx spatial gene signatures we derived, we found that genes for the *Normal Duct* signature mapped to the Ductal 1 cluster while the *LG-IPMN* signature was enriched in Ductal 2. Ductal 3 was enriched for the *Hyb-IPMN* gene signature while Ductal 4 shared transcripts with our *Advanced Neoplasia* signature (**Figure 4E)**. We then scored each ductal cell for our signatures and mapped each cell on a 3-way ternary plot to determine the relative enrichment of the *LG-IPMN, Hyb-IPMN,* and *Advanced Neoplasia* signature in each cell (**Figure 4F**). As expected, most cells from the side branch IPMN closely aligned with the *LG-IPMN* signature, consistent with its clinical pathological diagnosis. In contrast, main duct IPMN cells exhibited greater enrichment for the *Hyb-IPMN* and *Advanced Neoplasia* signatures. Tumor-derived cells showed the lowest enrichment for the *LG-IPMN* signature while displaying the highest enrichment for the *Advanced Neoplasia* signature **(Figure 4F)**. These data show that the *Hyb-IPMN* signature is distinctly present at the single cell level which further emphasizes shared features with malignant cells. We were encouraged by the finding that our side branch IPMN sample, which histologically contained only low-grade dysplasia, had a small but present population of cells enriched in the *Hyb-IPMN* signature, prompting us to investigate this finding further.

### An intermediary population of LG-IPMN with malignant transcriptional features is present in non-tumor bearing IPMN samples

As every sample in our spatial transcriptomic analysis was derived from tumor-bearing IPMNs encompassing the full spectrum of cell types from benign to malignant, one possible explanation is that the Hyb-IPMN population arises due to a field effect in tumor-affected pancreata, where benign cells near tumors are altered in response to tumor signaling. Notably, we have previously demonstrated that tissue adjacent to pancreatic tumors differs significantly from healthy tissue found in non-diseased pancreata[36]. We thus sought to analyze isolated LG-IPMN without associated cancer to determine if Hyb-IPMN is present in the absence of invasive disease. To answer this question, we queried a published external dataset that utilized the 10x Genomics Visium ST platform to characterize a cohort of patient tissues including non-tumor bearing LG-IPMN, non-tumor bearing HG-IPMN, and IPMN-derived PDAC [19].

Extracting spots with greater than 80% fraction of epithelial cells based on published Robust Cell Type Deconvolution scores, we mapped our GeoMx signatures onto these spots and found that while there was some degree of overlap, the *LG-IPMN, Hyb-IPMN,* and *Advanced Neoplasia* signatures were largely distinct **(Figure 5A-B)**. Our *LG-IPMN* signature was specifically enriched in LG-IPMN samples, while our *Hyb-IPMN* signature had some enrichment in a portion of LG-IPMN samples, with overall increasing enrichment in HG-IPMN and PDAC samples, respectively (**Figure 5B)**. Separating the samples on a patient-specific level, the *Hyb-IPMN* signature was found to have marked heterogeneous expression, both within each patient, and between patients (**Figure 5C)**. Extracting only the LG-IPMN samples, we annotated spots enriched for the *Hyb-IPMN* signature as Hyb-IPMN^HIGH^. Mapping genes from the *LG-IPMN* and *Hyb-IPMN* GeoMx signatures, we found near exclusive enrichment of *Hyb-IPMN* genes in the Hyb-IPMN^HIGH^ spots, while the *LG-IPMN* signature was expressed in both Hyb-IPMN^HIGH^ and Hyb-IPMN^LOW^ spots, but to a higher degree in the latter (**Figure 5D)**. We utilized spatial feature plots within the Visium pipeline to determine if spots enriched for the *Hyb-IPMN* signature were distinct from those enriched for the *LG-IPMN* signature and found that this was largely the case (**Figure 5E).**

**Figure 5:**
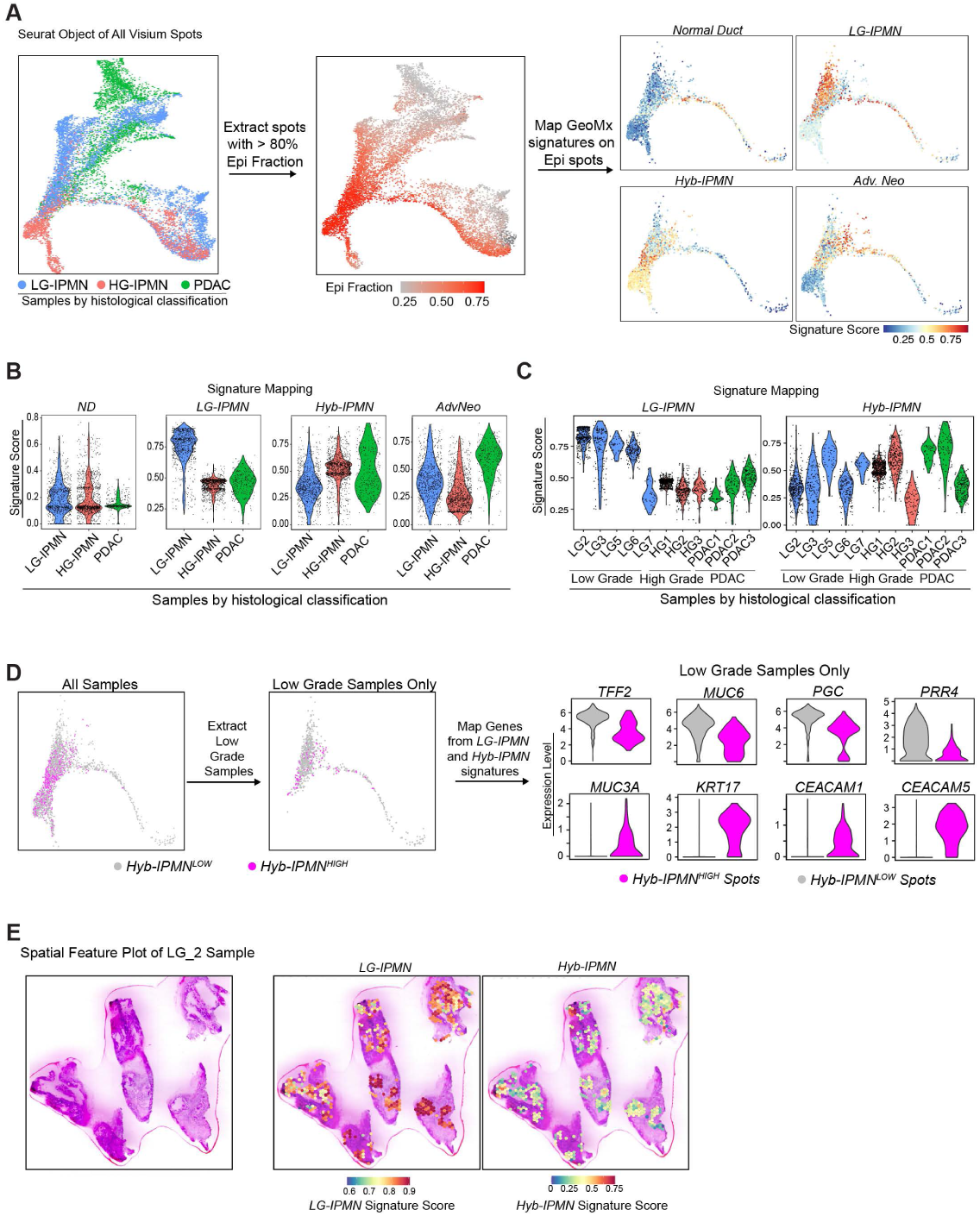
An intermediary population of LG-IPMN with malignant transcriptional features is present in non-tumor bearing IPMN samples. **A)** Left: UMAP plot of all spots from 7 LG-IPMN, 3 HG-IPMN, and 3 IPMN-associated PDAC samples. Epithelial spots (with epithelial fraction greater than 0.8) were extracted. Right: Scoring for signatures obtained from GeoMx spatial transcriptomics were mapped onto epithelial spots using AUcell score. **B)** Violin plots of the GeoMx signature scores plotted by histological classification. **C)** Violin plots of the GeoMx signature scores plotted by sample. **D)** Left: Scheme of workflow to compare epithelial spots with high enrichment of *Hyb-IPMN* signature vs. low enrichment of *Hyb-IPMN* signature. Right: Violin plots of genes from *LG-IPMN* signature (top) and *Hyb-IPMN* signature (bottom). E) Spatial feature plot of AUCell scoring of each spot of the *LG-IPMN* signature and *Hyb-IPMN* signature.

### Immunofluorescent staining of KRT17 and TFF2 in human IPMN tissues shows presence of a sub-population of KRT17 positive cells in LG-IPMN

With our bioinformatic analysis using NanoString GeoMx, single cell RNA sequencing, and Visium datasets, we have identified a broad spectrum of transcriptomes under the umbrella of IPMN epithelium that appears histologically low-grade, with a distinct subset harboring high-risk transcriptional features. To validate the existence of Hyb-IPMN as a distinct molecular entity from LG-IPMN at the protein level with a larger cohort, we next sought to directly visualize the presence of these cells *in situ* in a large cohort of patient pancreatic cysts utilizing 3 tissue microarrays (TMAs). Although the current WHO guidelines use a simplified two-tier grading system for IPMN, distinguishing only between low-grade and high-grade dysplasia[43], we opted to classify our samples using the earlier three-tier system (low-grade, intermediate-grade, and high-grade dysplasia). This approach allowed us to better correlate molecular changes with distinct histologic stages at a more granular level to parse the evolution of progression. Our cohort contained 51 LG-IPMN cores, 19 intermediate-grade IPMN cores, and 16 HG-IPMN cores. These tissues were stained with antibodies against TFF2 and KRT17, markers from the LG and Hyb-IPMN gene signature, respectively. We chose these particular markers as they are closely associated with the gastric foveolar (TFF2) and basal cell identity (KRT17); furthermore, loss of TFF2 and elevation of KRT17 expression have both been implicated in pancreatic tumorigenesis[25, 44, 45]. We found presence of KRT17 positive epithelial cells in HG-IPMN and IPMN-derived carcinoma, as well as in small patches of LG-IPMN (**Figure 6A)**. Stratifying by histologic grade, we observed that that TFF2 expression decreases as the degree of dysplasia increases (Figure 6B). With respect to KRT17, we observed that the majority of LG-IPMN cores showed minimal expression, with 85% of cores containing less than 5% KRT17-positive epithelial cells. The remaining LG-IPMN cores were heterogenous in KRT17 positivity, ranging from 5% to 30% of captured cyst epithelial cells (**Figure 6B)**. Stratifying TFF2 positivity and KRT17 positivity by IPMN subtype, we found that TFF2 expression was enriched in Gastric and Gastric/Intestinal (epithelium showing transition from gastric IPMN to intestinal IPMN) IPMN, while KRT17 positive cells were seen across all subtypes **(Supplemental Figure 5A-B)**. Interestingly, within the LG-IPMN cores, TFF2 positivity was inversely correlated with KRT17 positivity, with relatively sparse percentages of double positive cells **(Figure 6C)**, indicating that these markers define largely distinct cell populations. Of note, there was a significant number of epithelial cells that did not meet our threshold for positive staining, and thus were classified as double negative cells (**Supplemental Figure 5C)**. These data altogether confirm presence of KRT17 as a marker that is expressed in the cyst epithelial wall lining of HG-IPMN and carcinoma, and to a lesser degree in LG-IPMN, suggesting that there may be focal areas of LG-IPMN in pancreatic cysts that are at higher risk of progressing to advanced neoplasia.

**Figure 6:**
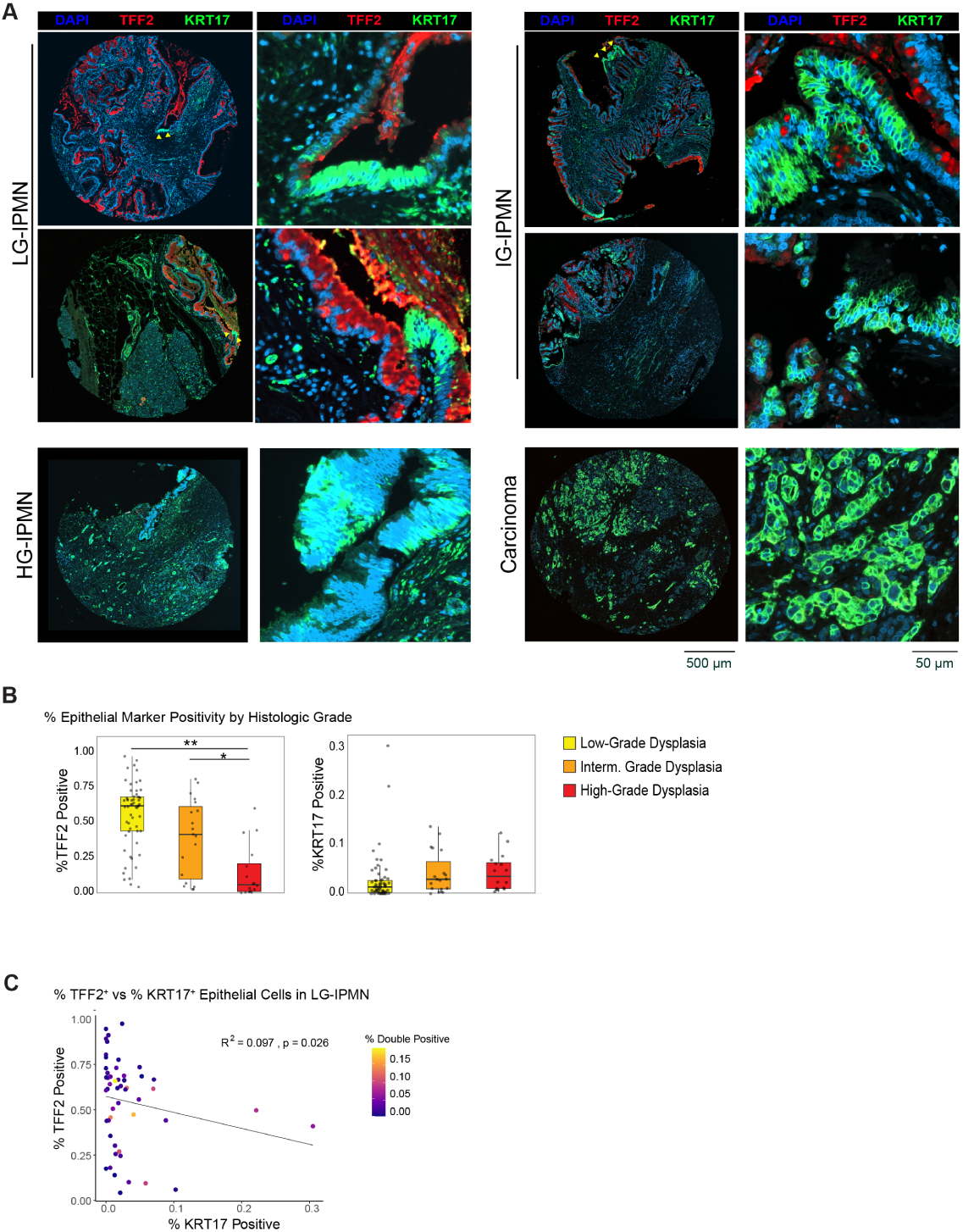
Immunofluorescent staining of KRT17 and TFF2 in human IPMN tissues shows prevalence of a sub-population of KRT17 positive cells in LG-IPMN. **A)** IPMN tissue of histologically defined LG-IPMN, IG-IPMN (intermediary grade IPMN), HG-IPMN, and IPMN-associated carcinoma stained for KRT17 (green), TFF2 (red), and DAPI (blue). Right-sided panels for each group represent a higher magnification area of each sample. Yellow arrows denote small patches of KRT17 positive cyst epithelium. **B)** Quantification of TFF2^+^ epithelial cells and KRT17^+^ epithelial cells as a fraction of total cyst epithelial cells, stratified by degree of dysplasia. **C)** Scatterplot of percent TFF2^+^ epithelial cells plotted against percent KRT17^+^ epithelial cells for each LG-IPMN core. Each point corresponds to a single core and is color-coded to indicate the proportion of double positive (KRT17^+^TFF2^+^) cyst epithelial cells within that core.

## Discussion

Here, we demonstrate the existence of a subpopulation of LG-IPMN epithelial cells that harbors high-risk transcriptional features overlapping with HG-IPMN and PDAC. We hypothesize that this subpopulation, referred to here as Hybrid-IPMN (Hyb-IPMN), represents a transitional state between indolent, low-grade disease and high-risk disease warranting intervention. These data are consistent with a prior IPMN single-cell RNA sequencing study that described 8.9% of LG-IPMN epithelial cells clustering with HG-IPMN epithelium [17]. We defined specific gene markers and signatures that may correspond to more aggressive disease in the Hyb-IPMN subgroup. In two ST datasets containing both IPMN with associated tumor and non-tumor bearing isolated IPMN samples, we demonstrate that there is a loss of expression of gastric genes such as TFF2, MUC6, and PGC and concomitant enrichment of genes associated with higher-risk disease, most notably KRT17, CLDN4, S100 proteins and CEA proteins prior to histologic progression. Finally, we show in a larger cohort of patient pancreatic cyst tissue that KRT17 expression is heterogeneous and prevalent in histologically low-grade IPMN, leading to the potential of its use in a risk stratification model of pancreatic cysts.

Interestingly, while initially the Hyb-IPMN entity appeared to exist on a transcriptomic continuum between LG-IPMN and HG-IPMN as suggested by other recent profiling studies[19, 34], we did find suggestion of a “branching” transitory point as Hyb-IPMN regions had markedly less gastric cell identity compared to LG-IPMN and HG-IPMN. This is consistent with a previous study focused on genetic alterations in IPMN progression which also found that IPMNs may progress to PDAC via sequential, branch-off, and do-novo evolutionary mechanisms[46]. These data points lend insight on real-world outcomes of patients with IPMN without worrisome features, as a small proportion will progress to malignancy, while most will not over the course of decades.

Regardless, this proportion does represent a clinical significant increased risk over the general population, with development of pancreatic cancer over a 10 year follow up at ∼6.6-8%, significantly increased over the general age-matched population[47, 48]. Interestingly, the risk of developing concomitant PDAC (not associated with the cyst), is increased in a subset of patients with larger cysts[48], pointing possibly to effects exerted on the microenvironment by IPMN, as progression to malignancy has been found to correlate with increased neutrophil infiltration and decreased plasma cell infiltration[19, 49]. This increased risk underscores the importance of identifying reliable biomarkers beyond imaging to help stratify the risk of malignant progression.

Our study has several limitations warranting further discussion. Most of the internal and external bioinformatic data was generated from resected IPMNs, and thus represent a single snapshot in time rather than serial examination during evolution of IPMN progression. Furthermore, resected IPMNs inevitably will represent a higher risk subset of pancreatic cysts, which is likely the reason that KRT17 positive cells were found to be prevalent, albeit in very small proportions, on staining validation of LG-IPMN cyst epithelium. In contrast to cross-sectional snapshots of IPMN, serial cyst fluid sampling is a potential avenue to correlate the development of marker expression with clinical outcome over time. However, this method has two important caveats; cyst fluid may not reflect the epithelial changes occurring in the cyst, and the only gold standard for evaluating for the presence of malignancy in a cyst is by careful pathological examination of the entire cyst lining, which can only be done in the setting of surgical resection. While there are other methods of evaluating the cyst epithelium endoscopically (via endoscopic cyst wall biopsy with microforceps and confocal microendoscopy[50, 51]) to allow for serial examination, these methods are limited in assessing the cyst wall in its entirety. Cyst wall biopsies, additionally, are associated with significant risk of bleeding and pancreatitis, and are thus not regularly performed in standard practice[52].

With regards to *in vivo* clinical models of IPMN, murine models that develop cystic lesions resembling IPMN, leveraging both KRAS and GNAS mutations, have been used to study the disease; however, they are limited in that they do not recapitulate the genetic heterogeneity[53, 54] or timing of progression of the disease[55]. Nevertheless, we are reassured by functional data showing that modulating gastric differentiation status can influence IPMN tumor progression in murine models [19, 56]. There also exists extensive literature on influence of genes in our Hyb-IPMN signature in PDAC outcomes in humans and modulation of PDAC aggressiveness *in vitro* and *in vivo* [25, 30, 31]. This study also has technical limitations in the spatial transcriptomic platforms used. Both platforms utilize ROIs or spots containing dozens to over a hundred cells. As the NanoString GeoMx ROIs were drawn based on epithelial histology and Visium spots have a fixed placement on the slide, these data points represent a mixed population of KRT17+/TFF2- Hyb-IPMN cells and KRT17-/TFF2+ LG-IPMN cells. Thus, it remains that our signatures may be further refined with better resolution methods. Finally, the genetic events promoting progression from LG-IPMN to Hyb-IPMN remain unclear, and one possibility worth exploring is whether genetic alterations precede transcriptomic changes in Hyb-IPMN, and whether there is a distinct mutational profile in Hyb-IPMN compared to LG-IPMN. Whatever the case may be, further integrated genomic and transcriptomic studies at single-cell and spatial resolution on KRT17-expressing cells low-grade IPMN will be revealing.

Recent studies have focused on cyst fluid or circulating DNA, protein, and RNA-based biomarker approaches and have identified promising targets for risk stratification in IPMN patients [53, 57–60]. The most significant advance has been the PancreaSeq test, which utilizes a next-generation sequencing approach to analyze cyst fluid obtained during endoscopic ultrasonography [53, 59]. PancreaSeq has been shown to be useful in stratifying cysts for malignant potential, with detection of a MAPK/GNAS mutation alongside alterations in TP53, SMAD4, CTNBB1, or mTOR exhibiting 88% sensitivity and 98% specificity for advanced neoplasia, warranting intervention. [53]. This test, while available clinically, is not yet widely implemented in standard practice. Further studies in characterizing the Hyb-IPMN population could help clarify how these genomic alterations translate phenotypically to a higher risk of malignancy, given that the underlying pathophysiology is not well-understood.

In summary, we demonstrate the presence of a high-risk subgroup within LG-IPMN epithelium that harbors malignant transcriptional features and altered cell identity, representing a transitional state between indolent and aggressive disease prior to histologic progression. We propose that this population may help to better understand the progression of IPMN and ultimately help to improve risk stratification of patients with pancreatic cysts, for which current clinical guidelines lack sufficient specificity to guide decision-making on surveillance versus intervention.

## Methods

### Patient Biospecimens

For single cell and spatial transcriptomics studies, medical chart review was used to screen for potential study patients with pancreatic cysts at the University of Michigan. *Endoscopic Fine Needle Biopsy:* Patients over the age of 18 years who had been referred for diagnostic endoscopic ultrasound of a pancreatic cyst were consented according to HUM00025339. Two extra passes using a 22 G SharkCore needle were taken for research after all specimens for clinical care had been obtained. *Surgical specimens:* Surgical specimens of IPMN tissue were obtained from patients referred for Whipple procedure or distal pancreatectomy according to IRB HUM00025339. Fresh specimens were procured by Tissue Procurement Services at the University of Michigan. Archival surgical FFPE specimens of pancreatic cysts seen at the University of Michigan from 2018 to 2022 were requested from Clinical Pathology. *Tissue Microarray*: Separately, three archival FFPE tissue microarrays (TMAs) were requested and obtained from University of Pittsburgh (IRB 19070069). Briefly, the TMAs were created from surgical resection material of consecutive pancreatic cyst patients seen at the University of Pittsburgh Medical Center (UPMC) between 2018 to 2023. For each surgical specimen, two 2-mm cores were obtained from a representative section of the pancreatic cyst to include both the epithelial and surrounding stromal component. Additionally, TMAs were created with aid of the TMA Grand Master^TM^ (3DHistotech) within the UPMC Hillman Translational Oncologic Pathology Services (TOPS).

### GeoMx Spatial Transcriptomics

Patient FFPE blocks containing gastric-subtype (sample 6132) or intestinal-subtype (samples 6276 and 6971) IPMN and associated carcinoma were chosen for spatial transcriptomics. Slides were prepared and processed at the University of Michigan Advanced Genomics Core according to manufacturer instructions utilizing the Whole Transcriptome Atlas (WTA) panel of genes and the Solid Tumor (TME) Morphology Kit. User-selected epithelial ROIs were segmented by PanCK expression to enrich for transcripts from epithelial populations, using PanCK+ for ductal ROIs and PanCK- for acinar. Quality control (QC) and normalization were performed according to the NanoString GeoMx recommended workflow. The following QC parameters were utilized to filter ROIs: >1000 reads, >80% trimmed, >80% stitched, > 80% aligned, > 50% sequencing saturation, >5 minNegative Count, < 3000 no template control count, > 100 nuclei, and area > 1000. Probes were then filtered to be detected in >5% of ROIs, and segments were filtered to have a gene detection rate of > 0.1. We used the limma package[61] batch correction function to regress out the batch effect, which showed the clustering of ROIs from the same cell type together, as we have done previously[36]. To define signatures of normal duct, LG-IPMN, Hyb-IPMN, and advanced neoplasia, we used a linear mixed model to perform DGE between each cell type and the rest of the ROIs. To score different cell identity signatures from the Panglao Dabatase[35] and Hallmark Genesets from MSigDB[62] for each ROI, we used GSVA[63].

### Single Cell RNA Sequencing

*Tissue processing:* For single cell processing, tissue was minced into 1mm^3^ pieces, then digested with 1 mg/mL collagenase P for 20-30 min at 37°C with gentle agitation. Digested tissue was rinsed three times with DMEM/1%BSA/10μM Y27632, then filtered through a 40μm mesh. Resulting live cells were submitted to the University of Michigan Advanced Genomics Core for 3’ single cell sequencing using the 10x Genomics Platform.

*Data Processing:* Samples were run using 50-cycle paired-end reads on the NovaSeq 6000 (Illumina) to a depth of 100,000 reads. Cell Ranger count version 6.0 was used with default settings, with an initially expected cell count of 10,000. The GRCh38 reference genome was used for alignment. Ambient RNA correction was done for each sample independently using SoupX[64]. We then performed Seurat’s recommended workflow[65]. Briefly, we removed low-quality cells with more than a 15% fraction of mitochondrial gene expression or less than 200 features (genes) detected. Expression was then log-normalized to the library size, and the top 2,000 highly variable features were extracted and scaled for downstream analysis. For sample integration, we used reciprocal PCA (rPCA). Count matrices were integrated to account for batch effects. We used UMAP dimension reduction to create two-dimensional embedding for the cells. For cell-type annotation, we used a panel of previously known markers for different cell types in pancreatic tissue to define different cell populations. Signature scoring of Panglao ductal cell, gastric foveolar, and basal cell identity were performed using AUCell[66]. Inferred Copy Number Variation analysis was performed on ductal epithelial clusters using InferCNV as provided at https://github.com/broadinstitute/inferCNV (inferCNV of the Trinity CTAT Project). Ductal epithelial cells from healthy donors were used as the reference cell type.

### Visium Spatial Transcriptomics

IPMN Visium post-processed and annotated data, as previously published[19], was downloaded from GEO accession GSE233254 in Seurat Object format. Epithelial spots were extracted by determination of epithelial fraction from Robust Cell Type Deconvolution[67] as greater than 0.8. Signature scoring of GeoMx-derived signatures was performed by AUCell Score[66]. The determination of Hyb-IPMN^LOW^ vs. Hyb-IPMN^HIGH^ spots was determined by the third quartile as a cut-off value utilizing the quantile function.

### Immunofluorescence

Archival FFPE pancreas tissue was sectioned at a thickness of 5-μm and mounted on charged slides. They were then baked at 60 degrees for 1 hour, deparaffinized, and dehydrated. They were then washed with deionized water and antigen retrieval was performed using a prewarmed steamer in sodium citrate buffer for 20 minutes. The TSA kit (Invitrogen) was used according to manufacturer’s instructions. Slides were then cooled to room temperature, washed with PBS, and incubated with 3% hydrogen peroxide for 15 minutes at room temperature. Slides were then incubated with blocking buffer for one hour at room temperature. Slides were then incubated overnight with primary antibody (rabbit anti-KRT17, 1:1000 dilution) at 4°C. Slides were then washed with PBS and incubated with anti-rabbit biotinylated secondary antibody from the TSA kit for one hour at room temperature. After additional PBS washes, slides were incubated with TSA conjugated to 488 for 8 minutes followed by stop solution for 5 minutes. They were then washes again with PBS and antigen retrieval, blocking, and primary antibody incubation (rabbit anti-TFF2, 1:500 dilution) were repeated as above. After overnight primary incubation, sections were washed with PBS and incubated with anti-rabbit 594 fluorescent secondary antibody (1:500 dilution) for one hour at room temperature. Slides were then washed with PBS, incubated with DAPI for 5 minutes, washed again with PBS, and mounted with Prolong Diamond Antifade.

### Microscopy and Image analysis

Tile-scanned fluorescent images were obtained using the Zeiss CellDiscoverer 7 microscope. Using QuPath 0.6.0, a random forest classifier to distinguish cyst epithelial cells from non-cyst epithelial cells was trained with pathologist annotation, followed by a thresholding classifier to detect marker positivity for KRT17 and TFF2. Percent positive cells were calculated as number of positive cyst epithelial cells divided by total number of cyst epithelial cells.

## Data Availability

Raw human data from the Carpenter, et. al. study[36, 68] are available at the National Institutes of Health (NIH) dbGaP database under the accession phs003229, with processed data available at NIH Gene Expression Omnibus database GSE229413. Additional unpublished sequencing files in this manuscript will be uploaded to new study accessions upon acceptance.

## Code Availability

All code used for this study will be made publicly available at the following GitHub upon acceptance.

**Please note that for ease of reviewing, we have made the unpublished transcriptomic data files and bioinformatics code available via a folder at the following link: https://www.dropbox.com/scl/fo/bp53i8gi6pvvytvd69d5w/ABAdtmzTCc9uqsoywvx8bA8?rlkey=bhssmthvgsbcleh6qa4l6vnla&st=swio2921&dl=0

password: Pancrea$

## Disclosures

AS reports the following disclosures: Apollo Endosurgery - Consultant, Advisory Board, Boston Scientific – Consultant, Olympus – Consultant, MicroTech – Consultant, GI Dynamics - Research / grant support, Fractyl - Consultant, Research / grant support. VS reports the following disclosures: Institutional research grant funding – Actuate Therapeutics, Boehringer Ingram, Bristol-Myers Squibb, Clovis, Elicio, Esanik, Exelixis, Fibrogen, Jazz, Ipsen, NCI, PanCAN, Cornerstone (previously Rafael), Relay, Repare, Servier, Transthera, Consultant – AstraZeneca, Autem, Amplity (previously Lynx Group), Delcath, Elevar, HistoSonics, Incyte, Ipsen, Jazz, Servier, Tallac, TransThera, Provision of equipment/supplies – Cornerstone (previously Rafael), Travel/lodging – Japanese Society of Clinical Oncology, BinayTara Foundation, Cholangiocarcinoma Foundation, HistoSonics, Lynx Group/Amplity, NCCN, Chinese Association of Pancreatology. The rest of the authors report no relevant disclosures.

Data transparency: Data will be made publicly available to other researchers upon publication of this manuscript.

## Author contributions

Jay Li: Conceptualization, Software, Formal analysis, Investigation, Data Curation, Writing Georgina Branch: Formal analysis

Justin Macchia: Formal analysis, Investigation Ahmed Elhossiny: Software

Nandini Arya: Formal analysis, Investigation Padma Kadiyala: Investigation

Nicole Peterson: Resources Julia Liang: Formal analysis Richard S. Kwon: Resources Jorge D. Machicado: Resources Erik-Jan Wamsteker: Resources Allison R. Schulman: Resources George M. Philips: Resources Jonathan Xia: Resources

Aatur Singhi: Resources

Jiayun M. Fang: Investigation, Resources, Writing – Review & Editing Vaibhav Sahai: Resources, Review & Editing

Filip Bednar: Resources – Writing – Review & Editing Timothy L. Frankel: Resources, Writing – Review & Editing

Marina Pasca Di Magliano: Resources, Writing – Review & Editing Jiaqi Shi: Investigation, Resources, Writing – Review & Editing

Eileen S. Carpenter: Conceptualization, Methodology, Software, Formal analysis, Investigation, Resources, Data Curation, Writing – Review & Editing, Supervision, Project administration, Funding acquisition

## Acknowledgements

We sincerely thank all patients who have generously provided tissue for this research. We thank D. Postiff from the University of Michigan Tissue Procurement Core for procurement of surgical samples. We thank T. Tamsen, M. Hogan and J. Opp from the University of Michigan Advanced Genomics Core for their assistance with single cell RNA sequencing and spatial transcriptomics. We thank Christopher Strayhorn for histology services. We acknowledge the Rogel and Blondy Center for Pancreatic Cancer for its support of this study. Work in Carpenter Lab was supported by the Department of Veterans Affairs BLRD Career Development Award CDA IK2BX005875, the American College of Gastroenterology Career Development Award, and the Cancer Discovery Award by the University of Michigan Rogel Cancer Center. The UPMC Hillman Translational Oncologic Pathology Services (TOPS) is supported by NCI 5P30CA047904. The funders had no role in study design, data collection and analysis, decision to publish or preparation of the manuscript.

## Abbreviations

Computed tomography (CT) Differential expression (DE) Differentially expressed gene (DEG) Gene set enrichment analysis (GSEA) Low-grade (LG)

High-grade (HG) High-risk (HR) Hybrid (Hyb)

Intraductal papillary mucinous neoplasm (IPMN) Magnetic resonance imaging (MRI)

Principle components analysis (PCA) Pancreatic ductal adenocarcinoma (PDAC) Region of interest (ROI)

Tumor microarray (TMA)

Uniform manifold approximation and projection (UMAP) Word count: 4030 words

Synopsis

We utilized a spatial transcriptomic approach to analyze gene expression profiles across the spectrum of IPMN disease. We identified a subpopulation of histologically low-grade IPMN harboring higher-risk malignant features and validated this population with immunostaining in patient tissues.

---

**Supplementary Figure 1.**
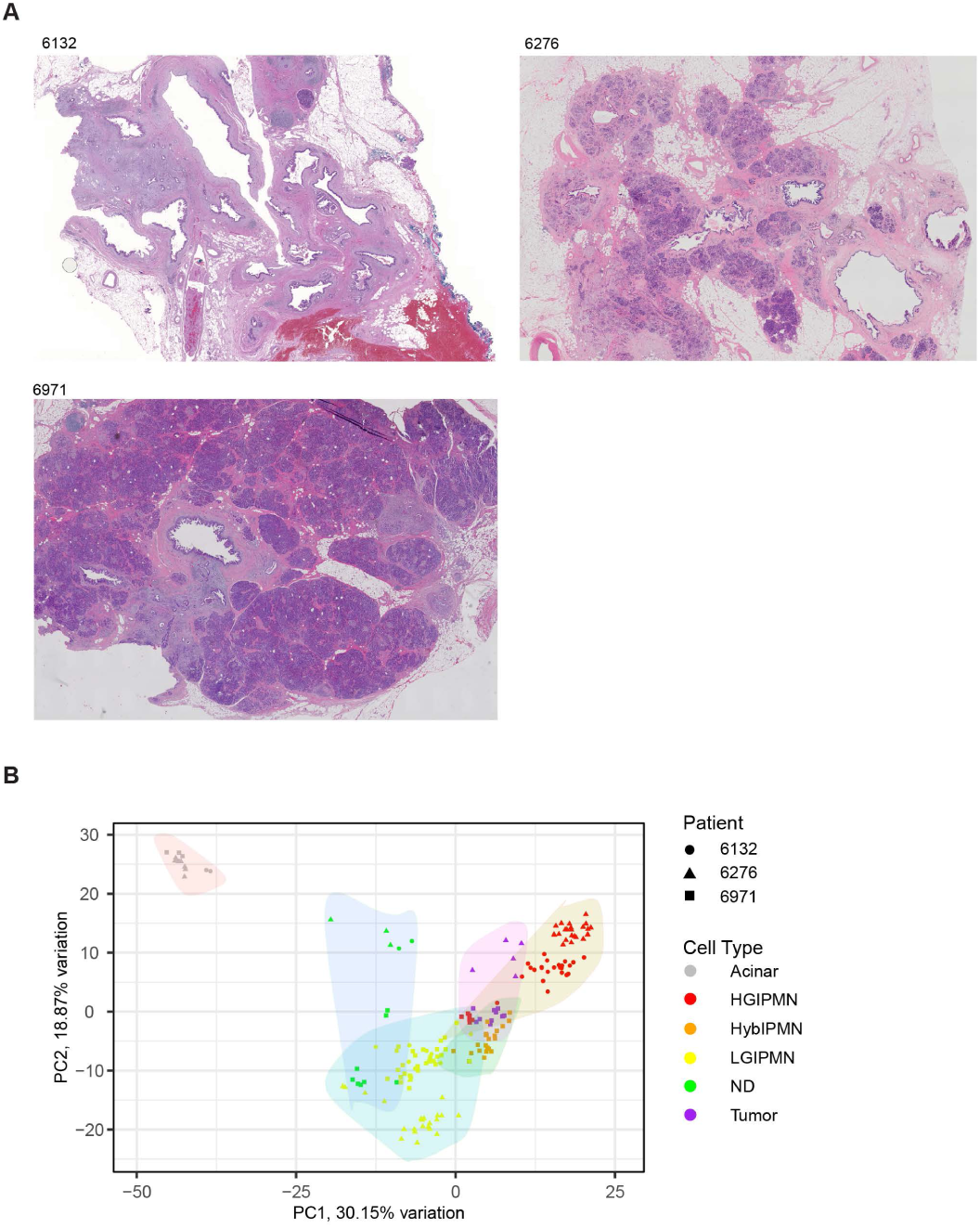
: Histology of IPMN tissue sections submitted for GeoMX Nanostring Spatial Transcriptomics. A) Low-magnification slide scans of serial FFPE H&E sections of the three IPMN samples submitted for spatial transcriptomics. B) PCA plot of epithelial segmented ROIs, marked by patient and cell type.

**Supplementary Figure 2.**
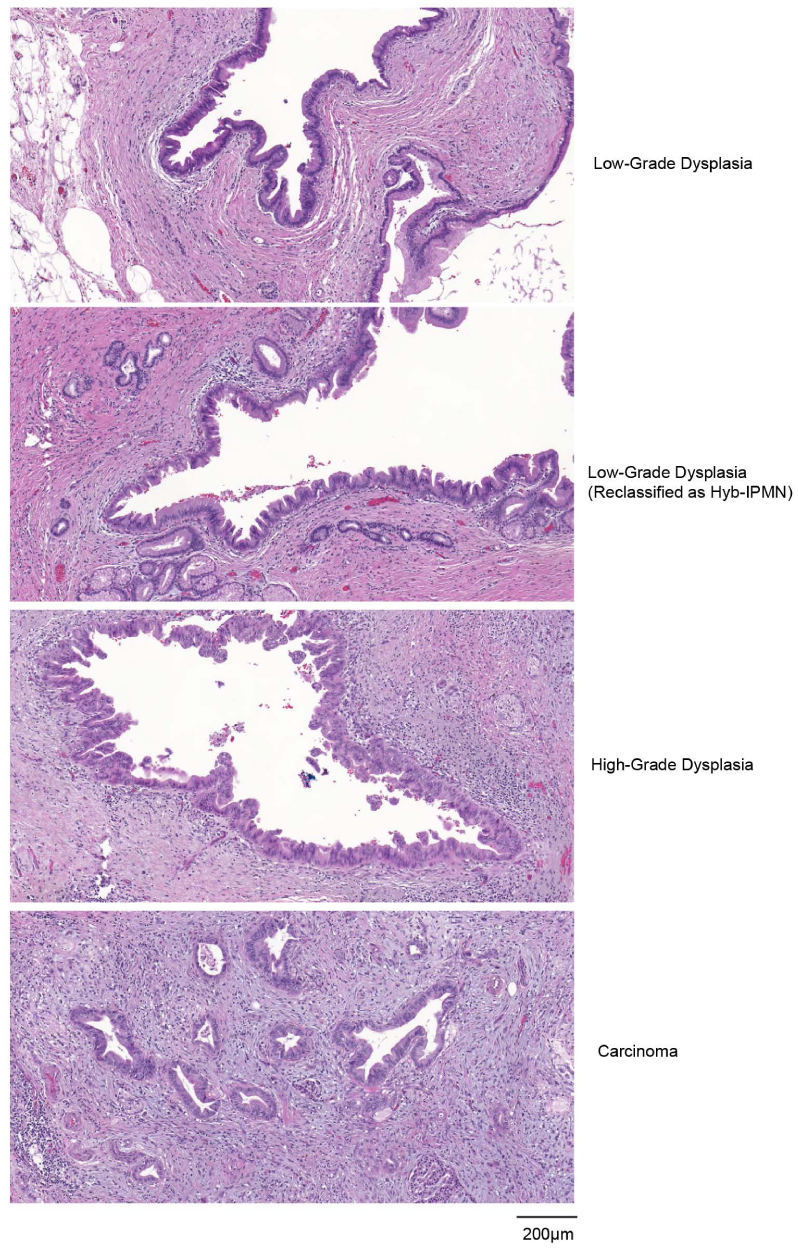
: Histology of Hyb-IPMN ROIs resemble that of LG-IPMN. High-magnification images of corresponding regions on serial H&E section of LG-IPMN, Hyb-IPMN, HG-IPMN, and carcinoma spatial transcriptomic ROIs.

**Supplementary Figure 3.**
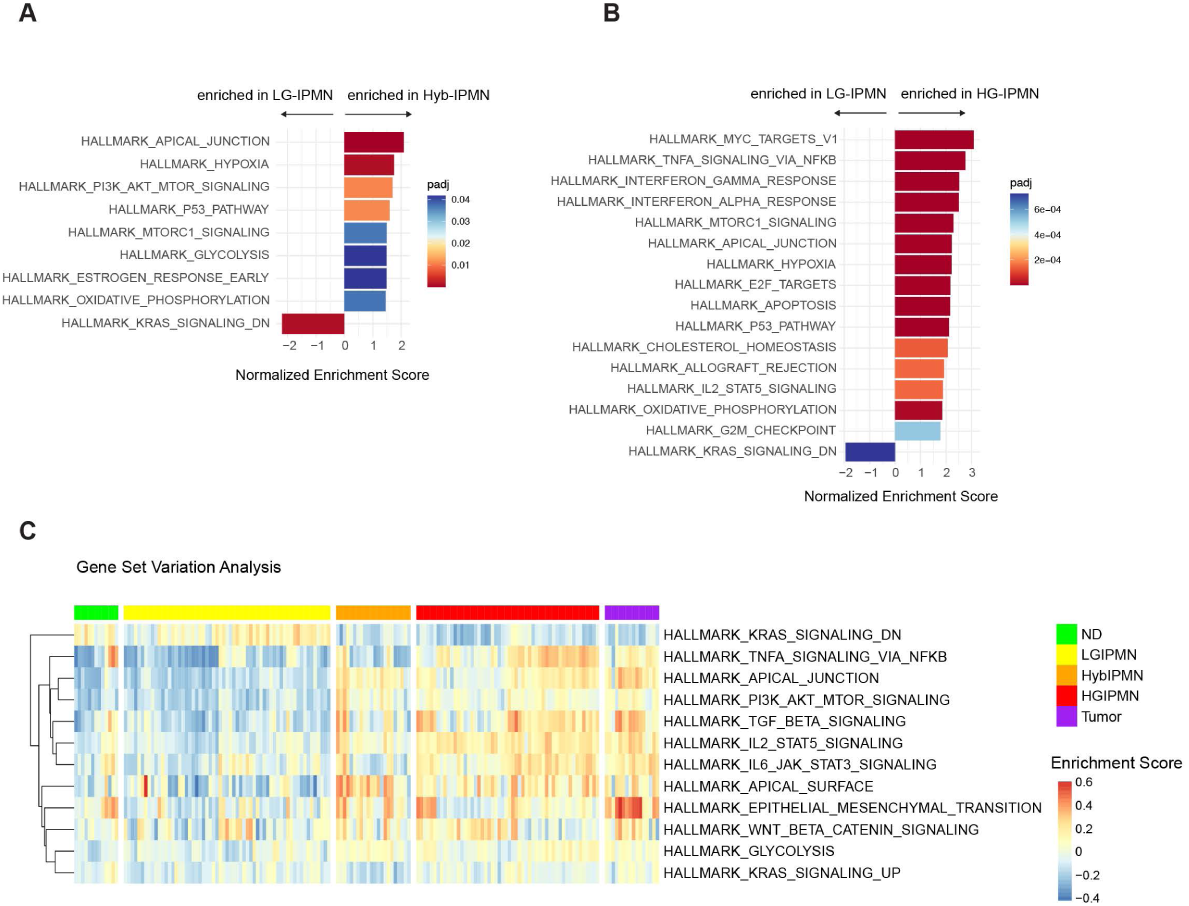
: Gene Set Variation Analysis shows shared and distinct features between Hyb-IPMN and HG-IPMN. A) Normalized enrichment score of significantly altered Hallmark genesets comparing Hyb-IPMN ROIs to LG-IPMN ROIs. B) Normalized enrichment score of significantly altered Hallmark genesets comparing HG-IPMN ROIs to LG-IPMN ROIs. C) Heatmap of normalized enrichment score across all ROIs, clustered by cell type.

**Supplementary Figure 4.**
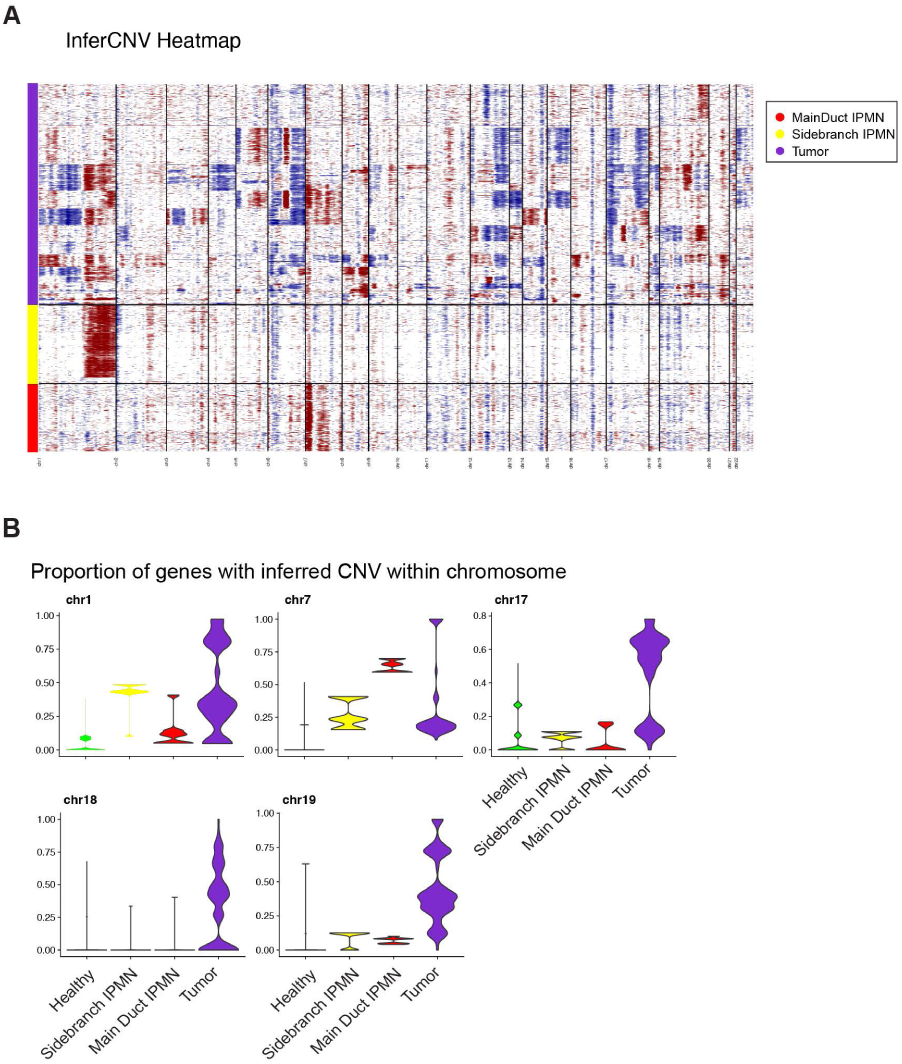
: IPMN have relatively sparse inferred copy number alterations. A) InferCNV heatmap of all Tumor, MainDuct IPMN, and Sidebranch IPMN ductal cells, using ductal cells from healthy donors as a reference control. Red color indicates inferred chromosomal amplification while blue color indicates inferred chromosomal deletion. B) Vlnplots of proportion of genes with inferred copy number variation within a given chromosome.

**Supplementary Figure 5.**
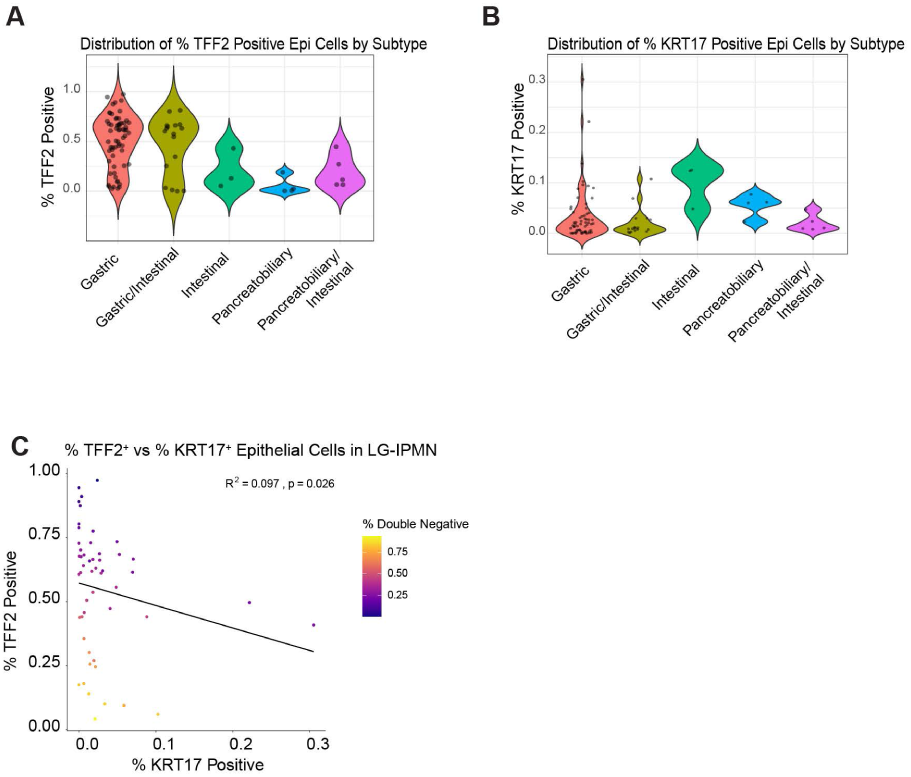
: KRT17 positive cells are seen across all IPMN histological subtypes A) VlnPlot of % TFF2 positive cells stratified by IPMN histological subtype. Gastric/Intestinal and Pancreatobiliary/Intestinal refers to cores that had mixing of both histological subtype within the cyst epithelium. Percent positive cells was calculated by dividing the number of positive cyst epithelial cells over the total number of cyst epithelial cells. B) VlnPlot of % KRT17 positive cells stratified by IPMN histological subtype. Gastric/Intestinal and Pancreatobiliary/Intestinal refers to cores that had mixing of both histological subtype within the cyst epithelium. Percent positive cells was calculated by dividing the number of positive cyst epithelial cells over the total number of cyst epithelial cells. C) Scatterplot of percent TFF2^+^ epithelial cells plotted against percent KRT17^+^ epithelial cells for each LG-IPMN core. Each point corresponds to a single core and is color-coded to indicate the proportion of double negative (KRT17^-^TFF2^-^) cyst epithelial cells within that core.

## References

[1] Siegel RL, Giaquinto AN, Jemal A. Cancer statistics, 2024. CA Cancer J Clin 2024;74(1):12–49.

[2] Rahib L, Wehner MR, Matrisian LM, Nead KT. Estimated Projection of US Cancer Incidence and Death to 2040. JAMA Netw Open 2021;4(4): e214708.

[3] Matthaei H, Schulick RD, Hruban RH, Maitra A. Cystic precursors to invasive pancreatic cancer. Nat Rev Gastroenterol Hepatol 2011;8(3):141–50.

[4] Vilela A, Quingalahua E, Vargas A, Hawa F, Shannon C, Carpenter ES, Shi J, Krishna SG, Lee UJ, Chalhoub JM, Machicado JD. Global Prevalence of Pancreatic Cystic Lesions in the General Population on Magnetic Resonance Imaging: A Systematic Review and Meta-analysis. Clin Gastroenterol Hepatol 2024;22(9):1798–809 e6.

[5] Wood LD, Adsay NV, Basturk O, Brosens LAA, Fukushima N, Hong SM, Kim SJ, Lee JW, Luchini C, Noe M, Pitman MB, Scarpa A, Singhi AD, Tanaka M, Furukawa T. Systematic review of challenging issues in pathology of intraductal papillary mucinous neoplasms. Pancreatology 2023;23(7):878–91.

[6] Furukawa T, Hatori T, Fujita I, Yamamoto M, Kobayashi M, Ohike N, Morohoshi T, Egawa S, Unno M, Takao S, Osako M, Yonezawa S, Mino-Kenudson M, Lauwers GY, Yamaguchi H, Ban S, Shimizu M. Prognostic relevance of morphological types of intraductal papillary mucinous neoplasms of the pancreas. Gut 2011;60(4):509–16.

[7] Kuboki Y, Shimizu K, Hatori T, Yamamoto M, Shibata N, Shiratori K, Furukawa T. Molecular biomarkers for progression of intraductal papillary mucinous neoplasm of the pancreas. Pancreas 2015;44(2):227–35.

[8] Ohtsuka T, Tomosugi T, Kimura R, Nakamura S, Miyasaka Y, Nakata K, Mori Y, Morita M, Torata N, Shindo K, Ohuchida K, Nakamura M. Clinical assessment of the GNAS mutation status in patients with intraductal papillary mucinous neoplasm of the pancreas. Surg Today 2019;49(11):887–93.

[9] Lee JH, Kim Y, Choi JW, Kim YS. KRAS, GNAS, and RNF43 mutations in intraductal papillary mucinous neoplasm of the pancreas: a meta-analysis. Springerplus 2016;5(1):1172.

[10] Shimizu T, Akita M, Sofue K, Toyama H, Itoh T, Fukumoto T, Zen Y. Pancreatobiliary-type intraductal papillary mucinous neoplasm of the pancreas may have 2 subtypes with distinct clinicopathologic and genetic features. Hum Pathol 2019;91:26–35.

[11] de la Fuente J, Chatterjee A, Lui J, Nehra AK, Bell MG, Lennon RJ, Kassmeyer BA, Graham RP, Nagayama H, Schulte PJ, Doering KA, Delgado AM, Vege SS, Chari ST, Takahashi N, Majumder S. Long-Term Outcomes and Risk of Pancreatic Cancer in Intraductal Papillary Mucinous Neoplasms. JAMA Netw Open 2023;6(10):e2337799.

[12] Tanaka M, Fernandez-Del Castillo C, Kamisawa T, Jang JY, Levy P, Ohtsuka T, Salvia R, Shimizu Y, Tada M, Wolfgang CL. Revisions of international consensus Fukuoka guidelines for the management of IPMN of the pancreas. Pancreatology 2017;17(5):738–53.

[13] European Study Group on Cystic Tumours of the P. European evidence-based guidelines on pancreatic cystic neoplasms. Gut 2018;67(5):789–804.

[14] van Huijgevoort NCM, Hoogenboom SAM, Lekkerkerker SJ, Busch OR, Del Chiaro M, Fockens P, Somers I, Verheij J, Voermans RP, Besselink MG, van Hooft JE. Diagnostic accuracy of the AGA, IAP, and European guidelines for detecting advanced neoplasia in intraductal papillary mucinous neoplasm/neoplasia. Pancreatology 2023;23(3):251–7.

[15] Dbouk M, Brewer Gutierrez OI, Lennon AM, Chuidian M, Shin EJ, Kamel IR, Fishman EK, He J, Burkhart RA, Wolfgang CL, Hruban RH, Goggins MG, Canto MI. Guidelines on management of pancreatic cysts detected in high-risk individuals: An evaluation of the 2017 Fukuoka guidelines and the 2020 International Cancer of the Pancreas Screening (CAPS) consortium statements. Pancreatology 2021;21(3):613–21.

[16] Frankel TL, LaFemina J, Bamboat ZM, D’Angelica MI, DeMatteo RP, Fong Y, Kingham TP, Jarnagin WR, Allen PJ. Dysplasia at the surgical margin is associated with recurrence after resection of non-invasive intraductal papillary mucinous neoplasms. HPB (Oxford) 2013;15(10):814–21.

[17] Bernard V, Semaan A, Huang J, San Lucas FA, Mulu FC, Stephens BM, Guerrero PA, Huang Y, Zhao J, Kamyabi N, Sen S, Scheet PA, Taniguchi CM, Kim MP, Tzeng CW, Katz MH, Singhi AD, Maitra A, Alvarez HA. Single-Cell Transcriptomics of Pancreatic Cancer Precursors Demonstrates Epithelial and Microenvironmental Heterogeneity as an Early Event in Neoplastic Progression. Clin Cancer Res 2019;25(7):2194–205.

[18] Matthaei H, Wylie D, Lloyd MB, Dal Molin M, Kemppainen J, Mayo SC, Wolfgang CL, Schulick RD, Langfield L, Andruss BF, Adai AT, Hruban RH, Szafranska-Schwarzbach AE, Maitra A. miRNA biomarkers in cyst fluid augment the diagnosis and management of pancreatic cysts. Clin Cancer Res 2012;18(17):4713–24.

[19] Sans M, Makino Y, Min J, Rajapakshe KI, Yip-Schneider M, Schmidt CM, Hurd MW, Burks JK, Gomez JA, Thege FI, Fahrmann JF, Wolff RA, Kim MP, Guerrero PA, Maitra A. Spatial Transcriptomics of Intraductal Papillary Mucinous Neoplasms of the Pancreas Identifies NKX6-2 as a Driver of Gastric Differentiation and Indolent Biological Potential. Cancer Discov 2023;13(8):1844–61.

[20] Iyer MK, Shi C, Eckhoff AM, Fletcher A, Nussbaum DP, Allen PJ. Digital spatial profiling of intraductal papillary mucinous neoplasms: Toward a molecular framework for risk stratification. Sci Adv 2023;9(11):eade4582.

[21] Heuer F, Sturmer R, Heuer J, Kalinski T, Lemke A, Meyer F, Hoffmann W. Different Forms of TFF2, A Lectin of the Human Gastric Mucus Barrier: In Vitro Binding Studies. Int J Mol Sci 2019;20(23).

[22] Hoffmann W. TFF2, a MUC6-binding lectin stabilizing the gastric mucus barrier and more (Review). Int J Oncol 2015;47(3):806–16.

[23] Terada T. An immunohistochemical study of primary signet-ring cell carcinoma of the stomach and colorectum: II. Expression of MUC1, MUC2, MUC5AC, and MUC6 in normal mucosa and in 42 cases. Int J Clin Exp Pathol 2013;6(4):613–21.

[24] Shen S, Jiang J, Yuan Y. Pepsinogen C expression, regulation and its relationship with cancer. Cancer Cell Int 2017;17:57.

[25] Li D, Ni XF, Tang H, Zhang J, Zheng C, Lin J, Wang C, Sun L, Chen B. KRT17 Functions as a Tumor Promoter and Regulates Proliferation, Migration and Invasion in Pancreatic Cancer via mTOR/S6k1 Pathway. Cancer Manag Res 2020;12:2087–95.

[26] Roa-Pena L, Leiton CV, Babu S, Pan CH, Vanner EA, Akalin A, Bandovic J, Moffitt RA, Shroyer KR, Escobar-Hoyos LF. Keratin 17 identifies the most lethal molecular subtype of pancreatic cancer. Sci Rep 2019;9(1):11239.

[27] Tsutsumi K, Sato N, Tanabe R, Mizumoto K, Morimatsu K, Kayashima T, Fujita H, Ohuchida K, Ohtsuka T, Takahata S, Nakamura M, Tanaka M. Claudin-4 expression predicts survival in pancreatic ductal adenocarcinoma. Ann Surg Oncol 2012;19 Suppl 3:S491–9.

[28] Gebauer F, Wicklein D, Horst J, Sundermann P, Maar H, Streichert T, Tachezy M, Izbicki JR, Bockhorn M, Schumacher U. Carcinoembryonic antigen-related cell adhesion molecules (CEACAM) 1, 5 and 6 as biomarkers in pancreatic cancer. PLoS One 2014;9(11):e113023.

[29] Al-Ismaeel Q, Neal CP, Al-Mahmoodi H, Almutairi Z, Al-Shamarti I, Straatman K, Jaunbocus N, Irvine A, Issa E, Moreman C, Dennison AR, Emre Sayan A, McDearmid J, Greaves P, Tulchinsky E, Kriajevska M. ZEB1 and IL-6/11-STAT3 signalling cooperate to define invasive potential of pancreatic cancer cells via differential regulation of the expression of S100 proteins. Br J Cancer 2019;121(1):65–75.

[30] Chen X, Liu X, Lang H, Zhang S, Luo Y, Zhang J. S100 calcium-binding protein A6 promotes epithelial-mesenchymal transition through beta-catenin in pancreatic cancer cell line. PLoS One 2015;10(3):e0121319.

[31] Li T, Ren T, Huang C, Li Y, Yang P, Che G, Luo L, Chen Y, Peng S, Lin Y, Zeng L. S100A16 induces epithelial-mesenchymal transition in human PDAC cells and is a new therapeutic target for pancreatic cancer treatment that synergizes with gemcitabine. Biochem Pharmacol 2021;189:114396.

[32] Moffitt RA, Marayati R, Flate EL, Volmar KE, Loeza SG, Hoadley KA, Rashid NU, Williams LA, Eaton SC, Chung AH, Smyla JK, Anderson JM, Kim HJ, Bentrem DJ, Talamonti MS, Iacobuzio-Donahue CA, Hollingsworth MA, Yeh JJ. Virtual microdissection identifies distinct tumor- and stroma-specific subtypes of pancreatic ductal adenocarcinoma. Nat Genet 2015;47(10):1168–78.

[33] Delgado-Coka LA, Roa-Pena L, Babu S, Horowitz M, Petricoin EF, 3rd, Matrisian LM, Blais EM, Marchenko N, Allard FD, Akalin A, Jiang W, Larson BK, Hendifar AE, Picozzi VJ, Choi M, Shroyer KR, Escobar-Hoyos LF. Keratin 17 is a prognostic and predictive biomarker in pancreatic ductal adenocarcinoma. Am J Clin Pathol 2024;162(3):314–26.

[34] Agostini A, Piro G, Inzani F, Quero G, Esposito A, Caggiano A, Priori L, Larghi A, Alfieri S, Casolino R, Scaglione G, Tondolo V, Cammarota G, Ianiro G, Corbo V, Biankin AV, Tortora G, Carbone C. Identification of spatially-resolved markers of malignant transformation in Intraductal Papillary Mucinous Neoplasms. Nat Commun 2024;15(1):2764.

[35] Franzen O, Gan LM, Bjorkegren JLM. PanglaoDB: a web server for exploration of mouse and human single-cell RNA sequencing data. Database (Oxford) 2019;2019.

[36] Carpenter ES, Elhossiny AM, Kadiyala P, Li J, McGue J, Griffith BD, Zhang Y, Edwards J, Nelson S, Lima F, Donahue KL, Du W, Bischoff AC, Alomari D, Watkoske HR, Mattea M, The S, Espinoza CE, Barrett M, Sonnenday CJ, Olden N, Chen CT, Peterson N, Gunchick V, Sahai V, Rao A, Bednar F, Shi J, Frankel TL, Pasca di Magliano M. Analysis of Donor Pancreata Defines the Transcriptomic Signature and Microenvironment of Early Neoplastic Lesions. Cancer Discov 2023;13(6):1324–45.

[37] Tanaka M, Fernandez-del Castillo C, Adsay V, Chari S, Falconi M, Jang JY, Kimura W, Levy P, Pitman MB, Schmidt CM, Shimizu M, Wolfgang CL, Yamaguchi K, Yamao K, International Association of P. International consensus guidelines 2012 for the management of IPMN and MCN of the pancreas. Pancreatology 2012;12(3):183–97.

[38] Steele NG, Carpenter ES, Kemp SB, Sirihorachai V, The S, Delrosario L, Lazarus J, Amir ED, Gunchick V, Espinoza C, Bell S, Harris L, Lima F, Irizarry-Negron V, Paglia D, Macchia J, Chu AKY, Schofield H, Wamsteker EJ, Kwon R, Schulman A, Prabhu A, Law R, Sondhi A, Yu J, Patel A, Donahue K, Nathan H, Cho C, Anderson MA, Sahai V, Lyssiotis CA, Zou W, Allen BL, Rao A, Crawford HC, Bednar F, Frankel TL, Pasca di Magliano M. Multimodal Mapping of the Tumor and Peripheral Blood Immune Landscape in Human Pancreatic Cancer. Nat Cancer 2020;1(11):1097–112.

[39] Eckhoff AM, Fletcher AA, Landa K, Iyer M, Nussbaum DP, Shi C, Nair SK, Allen PJ. Multidimensional Immunophenotyping of Intraductal Papillary Mucinous Neoplasms Reveals Novel T Cell and Macrophage Signature. Ann Surg Oncol 2022;29(12):7781–8.

[40] Hong SM, Vincent A, Kanda M, Leclerc J, Omura N, Borges M, Klein AP, Canto MI, Hruban RH, Goggins M. Genome-wide somatic copy number alterations in low-grade PanINs and IPMNs from individuals with a family history of pancreatic cancer. Clin Cancer Res 2012;18(16):4303–12.

[41] Fujikura K, Hosoda W, Felsenstein M, Song Q, Reiter JG, Zheng L, Beleva Guthrie V, Rincon N, Dal Molin M, Dudley J, Cohen JD, Wang P, Fischer CG, Braxton AM, Noe M, Jongepier M, Fernandez-Del Castillo C, Mino-Kenudson M, Schmidt CM, Yip-Schneider MT, Lawlor RT, Salvia R, Roberts NJ, Thompson ED, Karchin R, Lennon AM, Jiao Y, Wood LD. Multiregion whole-exome sequencing of intraductal papillary mucinous neoplasms reveals frequent somatic KLF4 mutations predominantly in low-grade regions. Gut 2021;70(5):928–39.

[42] Semaan A, Bernard V, Wong J, Makino Y, Swartzlander DB, Rajapakshe KI, Lee JJ, Officer A, Schmidt CM, Wu HH, Scaife CL, Affolter KE, Nachmanson D, Firpo MA, Yip-Schneider M, Lowy AM, Harismendy O, Sen S, Maitra A, Jakubek YA, Guerrero PA. Integrated Molecular Characterization of Intraductal Papillary Mucinous Neoplasms: An NCI Cancer Moonshot Precancer Atlas Pilot Project. Cancer Res Commun 2023;3(10):2062–73.

[43] Nagtegaal ID, Odze RD, Klimstra D, Paradis V, Rugge M, Schirmacher P, Washington KM, Carneiro F, Cree IA, Board WHOCoTE. The 2019 WHO classification of tumours of the digestive system. Histopathology 2020;76(2):182–8.

[44] Jiang Z, Wu F, Laise P, Takayuki T, Na F, Kim W, Kobayashi H, Chang W, Takahashi R, Valenti G, Sunagawa M, White RA, Macchini M, Renz BW, Middelhoff M, Hayakawa Y, Dubeykovskaya ZA, Tan X, Chu TH, Nagar K, Tailor Y, Belin BR, Anand A, Asfaha S, Finlayson MO, Iuga AC, Califano A, Wang TC. Tff2 defines transit-amplifying pancreatic acinar progenitors that lack regenerative potential and are protective against Kras-driven carcinogenesis. Cell Stem Cell 2023;30(8):1091–109 e7.

[45] Carpenter ES, Kadiyala P, Elhossiny AM, Kemp SB, Li J, Steele NG, Nicolle R, Nwosu ZC, Freeman J, Dai H, Paglia D, Du W, Donahue K, Morales J, Medina-Cabrera PI, Bonilla ME, Harris L, The S, Gunchick V, Peterson N, Brown K, Mattea M, Espinoza CE, McGue J, Kabala SM, Baliira RK, Renollet NM, Mooney AG, Liu J, Bhalla S, Farida JP, Ko C, Machicado JD, Kwon RS, Wamsteker EJ, Schulman A, Anderson MA, Law R, Prabhu A, Coulombe PA, Rao A, Frankel TL, Bednar F, Shi J, Sahai V, Pasca di Magliano M. KRT17High/CXCL8+ tumor cells display both classical and basal features and regulate myeloid infiltration in the pancreatic cancer microenvironment. Clin Cancer Res 2023.

[46] Omori Y, Ono Y, Tanino M, Karasaki H, Yamaguchi H, Furukawa T, Enomoto K, Ueda J, Sumi A, Katayama J, Muraki M, Taniue K, Takahashi K, Ambo Y, Shinohara T, Nishihara H, Sasajima J, Maguchi H, Mizukami Y, Okumura T, Tanaka S. Pathways of Progression From Intraductal Papillary Mucinous Neoplasm to Pancreatic Ductal Adenocarcinoma Based on Molecular Features. Gastroenterology 2019;156(3):647–61 e2.

[47] Oyama H, Tada M, Takagi K, Tateishi K, Hamada T, Nakai Y, Hakuta R, Ijichi H, Ishigaki K, Kanai S, Kogure H, Mizuno S, Saito K, Saito T, Sato T, Suzuki T, Takahara N, Morishita Y, Arita J, Hasegawa K, Tanaka M, Fukayama M, Koike K. Long-term Risk of Malignancy in Branch-Duct Intraductal Papillary Mucinous Neoplasms. Gastroenterology 2020;158(1):226–37 e5.

[48] Assawasirisin C, Fagenholz P, Qadan M, Hernandez-Barco Y, Aimprasittichai S, Kambadakone A, Mino-Kenudson M, Ike A, Chen SY, Sheng C, Brugge W, Warshaw AL, Lillemoe KD, Fernandez-Del Castillo C. Unraveling the Long-term Natural History of Branch Duct Intraductal Papillary Mucinous Neoplasm: Beyond 10 years. Ann Surg 2025;281(1):154–60.

[49] Carollo F, La Grutta G. [Effects of progesterone therapy in a case of probable epileptic-endocrinal syndrome]. Boll Soc Ital Biol Sper 1972;48(17):473–7.

[50] Machicado JD, Chao WL, Carlyn DE, Pan TY, Poland S, Alexander VL, Maloof TG, Dubay K, Ueltschi O, Middendorf DM, Jajeh MO, Vishwanath AB, Porter K, Hart PA, Papachristou GI, Cruz-Monserrate Z, Conwell DL, Krishna SG. High performance in risk stratification of intraductal papillary mucinous neoplasms by confocal laser endomicroscopy image analysis with convolutional neural networks (with video). Gastrointest Endosc 2021;94(1):78–87 e2.

[51] Samarasena J, Yu A, Lee D, Hashimoto R, Lu Y, Thieu D, Mai D, Lee J, Chang K. EUS-guided through-the-needle biopsy for pancreatic cystic lesions. VideoGIE 2019;4(9):436–9.

[52] Londono Castillo J, Ramai D, Crino SF, Facciorusso A. Experience of the moray micro forceps biopsy for pancreatic cystic lesions: lessons and insights from the MAUDE database. Expert Rev Gastroenterol Hepatol 2021;15(12):1345–7.

[53] Paniccia A, Polanco PM, Boone BA, Wald AI, McGrath K, Brand RE, Khalid A, Kubiliun N, O’Broin-Lennon AM, Park WG, Klapman J, Tharian B, Inamdar S, Fasanella K, Nasr J, Chennat J, Das R, DeWitt J, Easler JJ, Bick B, Singh H, Fairley KJ, Sarkaria S, Sawas T, Skef W, Slivka A, Tavakkoli A, Thakkar S, Kim V, Vanderveldt HD, Richardson A, Wallace MB, Brahmbhatt B, Engels M, Gabbert C, Dugum M, El-Dika S, Bhat Y, Ramrakhiani S, Bakis G, Rolshud D, Millspaugh G, Tielleman T, Schmidt C, Mansour J, Marsh W, Ongchin M, Centeno B, Monaco SE, Ohori NP, Lajara S, Thompson ED, Hruban RH, Bell PD, Smith K, Permuth JB, Vandenbussche C, Ernst W, Grupillo M, Kaya C, Hogg M, He J, Wolfgang CL, Lee KK, Zeh H, Zureikat A, Nikiforova MN, Singhi AD. Prospective, Multi-Institutional, Real-Time Next-Generation Sequencing of Pancreatic Cyst Fluid Reveals Diverse Genomic Alterations That Improve the Clinical Management of Pancreatic Cysts. Gastroenterology 2023;164(1):117–33 e7.

[54] Fischer CG, Beleva Guthrie V, Braxton AM, Zheng L, Wang P, Song Q, Griffin JF, Chianchiano PE, Hosoda W, Niknafs N, Springer S, Dal Molin M, Masica D, Scharpf RB, Thompson ED, He J, Wolfgang CL, Hruban RH, Roberts NJ, Lennon AM, Jiao Y, Karchin R, Wood LD. Intraductal Papillary Mucinous Neoplasms Arise From Multiple Independent Clones, Each With Distinct Mutations. Gastroenterology 2019;157(4):1123–37 e22.

[55] Makino Y, Rajapakshe KI, Chellakkan Selvanesan B, Okumura T, Date K, Dutta P, Abou-Elkacem L, Sagara A, Min J, Sans M, Yee N, Siemann MJ, Enriquez J, Smith P, Bhattacharya P, Kim M, Dede M, Hart T, Maitra A, Thege FI. Metabolic reprogramming by mutant GNAS creates an actionable dependency in intraductal papillary mucinous neoplasms of the pancreas. Gut 2024;74(1):75–88.

[56] Yamaguchi J, Mino-Kenudson M, Liss AS, Chowdhury S, Wang TC, Fernandez-Del Castillo C, Lillemoe KD, Warshaw AL, Thayer SP. Loss of Trefoil Factor 2 From Pancreatic Duct Glands Promotes Formation of Intraductal Papillary Mucinous Neoplasms in Mice. Gastroenterology 2016;151(6):1232–44 e10.

[57] Yang KS, Ciprani D, O’Shea A, Liss AS, Yang R, Fletcher-Mercaldo S, Mino-Kenudson M, Fernandez-Del Castillo C, Weissleder R. Extracellular Vesicle Analysis Allows for Identification of Invasive IPMN. Gastroenterology 2021;160(4):1345–58 e11.

[58] Do M, Kim H, Shin D, Park J, Kim H, Han Y, Jang JY, Kim Y. Marker Identification of the Grade of Dysplasia of Intraductal Papillary Mucinous Neoplasm in Pancreatic Cyst Fluid by Quantitative Proteomic Profiling. Cancers (Basel) 2020;12(9).

[59] Singhi AD, McGrath K, Brand RE, Khalid A, Zeh HJ, Chennat JS, Fasanella KE, Papachristou GI, Slivka A, Bartlett DL, Dasyam AK, Hogg M, Lee KK, Marsh JW, Monaco SE, Ohori NP, Pingpank JF, Tsung A, Zureikat AH, Wald AI, Nikiforova MN. Preoperative next-generation sequencing of pancreatic cyst fluid is highly accurate in cyst classification and detection of advanced neoplasia. Gut 2018;67(12):2131–41.

[60] Yip-Schneider MT, Carr RA, Wu H, Schmidt CM. Prostaglandin E(2): A Pancreatic Fluid Biomarker of Intraductal Papillary Mucinous Neoplasm Dysplasia. J Am Coll Surg 2017;225(4):481–7.

[61] Ritchie ME, Phipson B, Wu D, Hu Y, Law CW, Shi W, Smyth GK. limma powers differential expression analyses for RNA-sequencing and microarray studies. Nucleic Acids Res 2015;43(7):e47.

[62] Liberzon A, Birger C, Thorvaldsdottir H, Ghandi M, Mesirov JP, Tamayo P. The Molecular Signatures Database (MSigDB) hallmark gene set collection. Cell Syst 2015;1(6):417–25.

[63] Hanzelmann S, Castelo R, Guinney J. GSVA: gene set variation analysis for microarray and RNA-seq data. BMC Bioinformatics 2013;14:7.

[64] Young MD, Behjati S. SoupX removes ambient RNA contamination from droplet-based single-cell RNA sequencing data. Gigascience 2020;9(12).

[65] Hao Y, Hao S, Andersen-Nissen E, Mauck WM, 3rd, Zheng S, Butler A, Lee MJ, Wilk AJ, Darby C, Zager M, Hoffman P, Stoeckius M, Papalexi E, Mimitou EP, Jain J, Srivastava A, Stuart T, Fleming LM, Yeung B, Rogers AJ, McElrath JM, Blish CA, Gottardo R, Smibert P, Satija R. Integrated analysis of multimodal single-cell data. Cell 2021;184(13):3573–87 e29.

[66] Aibar S, Gonzalez-Blas CB, Moerman T, Huynh-Thu VA, Imrichova H, Hulselmans G, Rambow F, Marine JC, Geurts P, Aerts J, van den Oord J, Atak ZK, Wouters J, Aerts S. SCENIC: single-cell regulatory network inference and clustering. Nat Methods 2017;14(11):1083–6.

[67] Cable DM, Murray E, Zou LS, Goeva A, Macosko EZ, Chen F, Irizarry RA. Robust decomposition of cell type mixtures in spatial transcriptomics. Nat Biotechnol 2022;40(4):517–26.

[68] Steele NG, Carpenter ES, Kemp SB, Sirihorachai VR, The S, Delrosario L, Lazarus J, Amir E-aD, Gunchick V, Espinoza C, Bell S, Harris L, Lima F, Irizarry-Negron V, Paglia D, Macchia J, Chu AKY, Schofield H, Wamsteker E-J, Kwon R, Schulman A, Prabhu A, Law R, Sondhi A, Yu J, Patel A, Donahue K, Nathan H, Cho C, Anderson MA, Sahai V, Lyssiotis CA, Zou W, Allen BL, Rao A, Crawford HC, Bednar F, Frankel TL, Pasca di Magliano M. Multimodal mapping of the tumor and peripheral blood immune landscape in human pancreatic cancer. Nature Cancer 2020;1(11):1097–112.

